# Multimodal dynamics control activity of a glial glutamate transporter

**DOI:** 10.64898/2026.05.29.728845

**Authors:** Qianyi Wu, Didar Ciftci, Juan Carlos Canul Tec, Nicolas Reyes, Gabriel Glenn Gregorio, Yun Huang, Olga Boudker

**Affiliations:** Department of Biochemistry & Biophysics, Weill Cornell Medicine, 1300 York Ave, New York, NY, 10065, USA; Howard Hughes Medical Institute, Weill Cornell Medicine, New York, NY, USA; Howard Hughes Medical Institute, Department of Chemistry and Chemical Biology, Department of Physics, Harvard University, Cambridge, MA 02138, USA; Fundamental Microbiology and Pathogenicity Unit, CNRS, Université de Bordeaux, IECB, Bordeaux, France; Sanavia Oncology, New York, NY, 10016, USA

## Abstract

Membrane transporters move polar solutes across lipid bilayers to regulate cellular metabolism, signaling, and drug distribution. These proteins operate via an alternating-access mechanism, cycling between extracellular-, intermediate-, and intracellular-facing conformations. The human excitatory amino acid transporter 1 (EAAT1) protects neurons from excitotoxic damage by mediating the uptake of glutamate and aspartate into glial cells. Defects in EAAT1 function result in numerous pathologies, including epilepsy and ataxia, suggesting that positive modulation of these transporters might ameliorate glutamate neurotoxicity. However, developing EAAT1 activators requires understanding the timing of conformational changes, which remain largely unexplored. Here, we establish an experimental platform that combines single-molecule Förster resonance energy transfer (smFRET) to monitor real-time conformational dynamics, single-transporter activity assays to correlate dynamics with function, and cryogenic electron microscopy (cryoEM) to visualize discrete conformations at high resolution. This platform enables detection of Ångstrom-scale movements of single transporter molecules in real time, revealing that EAAT1 intersperses rapid conformational dynamics with long pauses. Slow and fast dynamics can be modulated by substrates, membrane composition, and mutations, and are correlated with the enrichment of specific structural states. We leverage this platform to investigate an EAAT1 mutation associated with severe episodic ataxia and show that it inhibits transport by stabilizing a paused cytoplasm-facing conformation. These results identify multimodal dynamics as an intrinsic, regulatable feature of EAAT1 function and, therefore, a potential therapeutic target. Henceforth, our integrated platform will facilitate investigations of other regulatory factors, including the effects of small-molecule and lipid modulators on the transport cycle.

## Main

Most proteins transition between multiple structural conformations, and the rates of these transitions determine functional outcomes and influence cellular physiology. Members of the solute carrier family – integral membrane proteins that transport a wide range of substrates – exemplify this paradigm. By switching between outward-facing, intermediate, and inward-facing states (OFS, intS, and IFS, respectively), these transporters capture substrates on one side of the membrane and release them on the other (**ED Fig. 1a**). Although high-resolution structures have provided static snapshots of these states for several transporters, it remains unknown how transitions between them occur and how their dynamics determine transport activity.

Astroglial excitatory amino acid transporters, EAAT1 and EAAT2, support neurotransmission by rapidly removing glutamate (Glu) and aspartate (Asp) from the synaptic cleft. They move substrates against their concentration gradients by coupling uptake to the energetically favorable influx of three Na^+^ and one H^+^, and the efflux of one K^+ 1^. EAATs assemble into homotrimers and function via an elevator mechanism. The relatively static scaffold domains of each protomer mediate trimerization, whereas the transport domains, harboring substrate- and ion-binding sites, undergo ∼10-15 Å movements across the bilayer (**Fig. 1a**). EAATs also mediate thermodynamically uncoupled Cl^-^ currents ^2,3^ that are gated by substrate binding and modulate cell excitability ^4–6^. Because EAATs prevent excitotoxicity, their dysfunction contributes to a plethora of pathologies ^7–11^, including neuropathic pain, epilepsy, neurodegeneration, episodic ataxia, and addictive behavior ^12–20^. Thus, EAATs are promising drug targets, especially for positive modulators that could ameliorate Glu neurotoxicity ^20– 25^. However, since potentiating their function requires accelerating the conformational changes underlying substrate transport, understanding these transitions with sufficient temporal and spatial resolution is essential. Currently, there are no approaches to dissect transporters at this level of detail, hindering investigations of pathogenic mutations and the effects of environmental and pharmacological factors.

**Figure 1.**
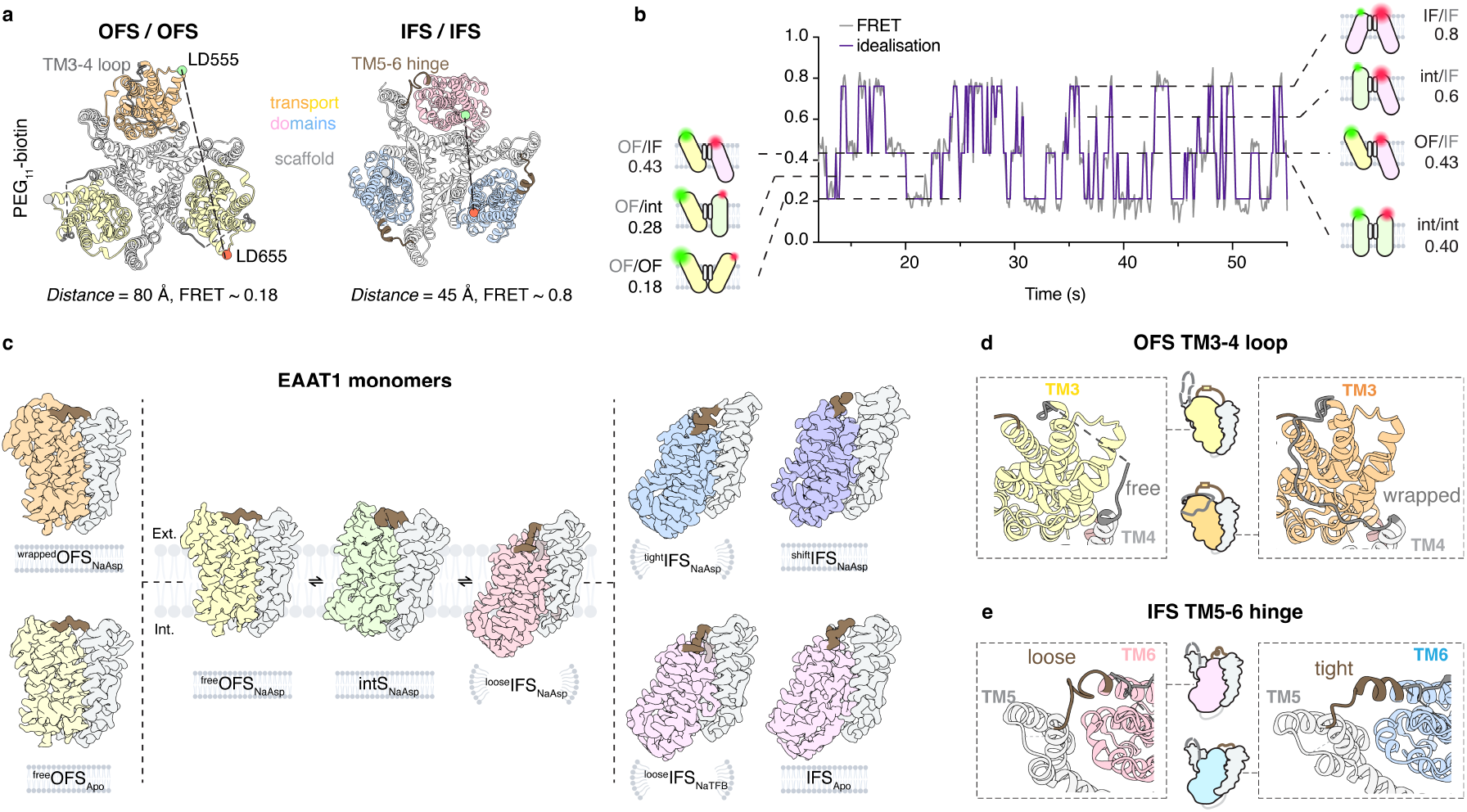
Structural snapshots enable smFRET state assignment. **a**, Cartoon representation of trimeric EAAT1_fret_ models with all protomers in either the OFS or IFS. Transport domain color scheme (here and throughout): ^free^OFS, light yellow; ^wrapped^OFS, orange; intS, green; ^loose^IFS, pink; ^tight^IFS, blue; ^shift^IFS, purple. Scaffold domains shown in white and extracellular hinges in gray (TM3-4) and brown (TM5-6). Labeling sites indicated as spheres: donor dye (LD555), green; acceptor dye (LD655), red; PEG_11_-biotin, silver. Distances and expected FRET efficiencies given below the structures. **b**, Example smFRET experiment: raw data, gray; idealized trace, purple. Dashed lines represent four distinct FRET states due to seven combinations of protomer pairs in outward-facing (OF), intermediate (int), and inward-facing (IF) configurations. Cartoon representations of protomer pairs and expected FRET efficiencies shown at left and right. **c**, cryoEM maps of individual protomers along the envisioned transport process. Cartoon micelles and bilayers around the state names indicate detergent or nanodiscs sample preparation, respectively. Detergent molecule under the TM5-6 hinge in ^loose^IFS highlighted in dusty pink. **d-e**, Close-up views of alternative hinge configurations: TM3-4 in the OFS (**d**); TM5-6 in the IFS (**e**). Schematics of EAAT1_fret_ protomers in each hinge configuration shown alongside.

To bridge this gap, we combined smFRET microscopy, single-transporter activity assays, and single-particle cryoEM to correlate Ångstrom-scale molecular motions with the activity of individual transporters *in situ* and in real time ^26–30^ and to uncover their structural underpinnings. We show that transport domains undergo stepwise movements along their scaffold domains, from the OFS through an intS to the IFS. Elevator dynamics are fast in the presence of Na^+^ and substrate or K^+^ alone, but not with Na^+^ alone, as expected for an ion-coupled transporter. Surprisingly, EAAT1 operates in multiple modes, alternating between rapid activity and periods with prolonged pauses. Membrane composition and mutations influence the dynamics and the balance between the modes. When active, protomers in EAAT1 trimers move independently; however, they preferentially pause in symmetric OF or IF states, suggesting inter-protomer communication. CryoEM structures reveal substates of both the OFS and IFS that differ in scaffold domain architecture, inter-domain hinges, and lipid interactions. Enrichment of these substates, achieved by rigidifying the hinges and scaffold domains, correlates with increased pausing. We leveraged these insights and the integrated platform to investigate the pathophysiological mechanism underlying the episodic ataxia-associated P290R mutation ^14^. We found that structural changes in the mutant scaffold lead to pronounced IFS pauses, resulting in significantly reduced Glu uptake and helping to explain increased uncoupled Cl^-^ currents, characteristic of this mutant.

We propose that EAAT1 can enter off-cycle kinetic traps, causing functional pauses that slow substrate uptake and alter Cl^-^ conductance. This suggests that modulation of EAAT1 can be achieved by affecting the transport cycle itself or by shifting the balance between activity and pausing. Our platform enables investigation of transporter regulation in cells, the impact of clinically relevant mutations, and pharmacological potentiation of activity.

### smFRET and cryoEM define EAAT1’s transport cycle

To detect movements of the EAAT1 transport domain in real time using smFRET, we used a thermally stabilized variant that is 95% identical to EAAT1 (**ED Fig. 1**). Unlike wild-type transporters, it can be purified in detergent solution and reconstituted into lipid vesicles without losing activity ^31^. We also introduced a cysteine mutation, N422C, into an extracellular loop of the transport domain to enable labeling with biotin and fluorescent reagents (**Fig. 1a, ED Fig. 1b-c, 2a**). The resulting construct (WT EAAT1_fret_ for brevity) was incubated with a mixture of maleimide-activated biotin-PEG_11_ and donor (LD555) and acceptor (LD655) fluorophores, yielding specific labeling with 57.0 ± 0.8% and 57.1 ± 1.5% efficiency for LD555 and LD655, respectively (**ED Fig. 2b**,**c**). Labeled proteins were reconstituted into liposomes at a low protein-to-lipid ratio to maximize the proportion of vesicles with single EAAT1_fret_ trimers. Proteoliposomes containing EAAT1_fret_, oriented with extracellular domains facing outward, were immobilized via a biotin-neutravidin bridge in microfluidic chambers for fluorescence imaging ^26,27^. Only smFRET recordings corresponding to single EAAT1_fret_ trimers containing donor and acceptor fluorophores were selected for analysis (see *Methods*).

smFRET between LD555- and LD655-labeled protomers showed transitions between multiple states (FRET values between ∼0.18 and 0.8; **Fig. 1b**). To assign these FRET states to specific conformations, we used cryoEM to image EAAT1_fret_ in detergent and nanodiscs under active (with 200 mM Na^+^ and 1 mM Asp or with 200 mM K^+^) and inhibited (with 200 mM Na^+^ and 400 µM blocker TFB-TBOA) conditions (**Fig. 1c**). Refinement with imposed C3 symmetry, followed by symmetry expansion and 3D classification, resolved conformational states for individual protomers (**ED Figs. 3-5** and **ED Table 1**). In K^+^, we observed only *apo* (K^+^-free) protomers in the OFS and IFS, consistent with previous reports that K^+^-bound states are too transient to detect ^32^. In the presence of Na^+^ and TFB-TBOA, protomers were in an IFS with an open substrate-binding site, resembling Na^+^-bound transporters after substrate release ^33,34^. In Na^+^/Asp, the protomers adopted three primary conformations (OFS, intS, and IFS), with substates within OFS and IFS. These conformational substates exhibited differences in the hinges and loops connecting the mobile transport domain to the static scaffold domain (**ED Fig. 1a**,**b**), particularly between transmembrane helices (TMs) 3 and 4, and 5 and 6. OFS protomers adopted “wrapped” and “free” substates, in which the extracellular TM3-4 loop wraps around the transport domain or is unstructured, respectively (**Fig. 1d**). IFS protomers populated “tight” and “loose” substates (**Fig. 1e**), with a helical or unstructured extracellular TM5-6 hinge, respectively.

We assigned three main FRET states for the fluorescently labeled protomer pairs (∼0.2, 0.4, and 0.8) based on inter-protomer C422-C422 distances in our structural models (**ED Table 2**) to OF/OF, OF/IF, and IF/IF, respectively. Notably, the OFS and IFS substates cannot be distinguished in smFRET because their C422-C422 distances are similar. intS forms OF/int, int/int, and IF/int configurations, which are minor because intS is a transient state. IF/int produces a distinct ∼0.6 FRET state, whereas OF/int (∼0.3) and int/int (∼0.4) overlap with OF/OF and OF/IF states (**Fig. 1b, ED Table 2**). Since our cryoEM structures captured most of the known states of the transport cycle (**Fig. 1c; ED Fig. 6**), we assigned all observed FRET states to structural conformations (**Fig. 1b; ED Table 2**), enabling observation of Ångstrom-scale structural changes with 100 msec temporal resolution.

### Substrates regulate dynamics and ion coupling

To concentrate substrates up to a million-fold within glial cells, EAAT1 must adhere to fundamental thermodynamic principles of ion-coupled transport: substrate and symported Na^+^ are moved together, not separately, and antiported K^+^ is required to reset the transporter to OFS to complete the cycle (**Fig. 2a**). To test these predictions with smFRET experiments, we configured substrate and ion exchange conditions using symmetric internal and external proteoliposome buffers. Since every step of the transport cycle is reversible, we expect to observe back-and-forth elevator transitions with symmetric Na^+^/Asp, during which Asp and symported Na^+^ ions are moved across the membrane in exchange for another Asp and Na^+^ ions (**Fig. 2a**). Similarly, K^+^ ions can be exchanged under symmetric conditions. Conversely, in a Na^+^ only buffer, or without any alkali ions, EAAT1 should be unable to exhibit elevator dynamics. After assigning structural conformations to the observed smFRET states, we experimentally tested these predictions.

**Figure 2.**
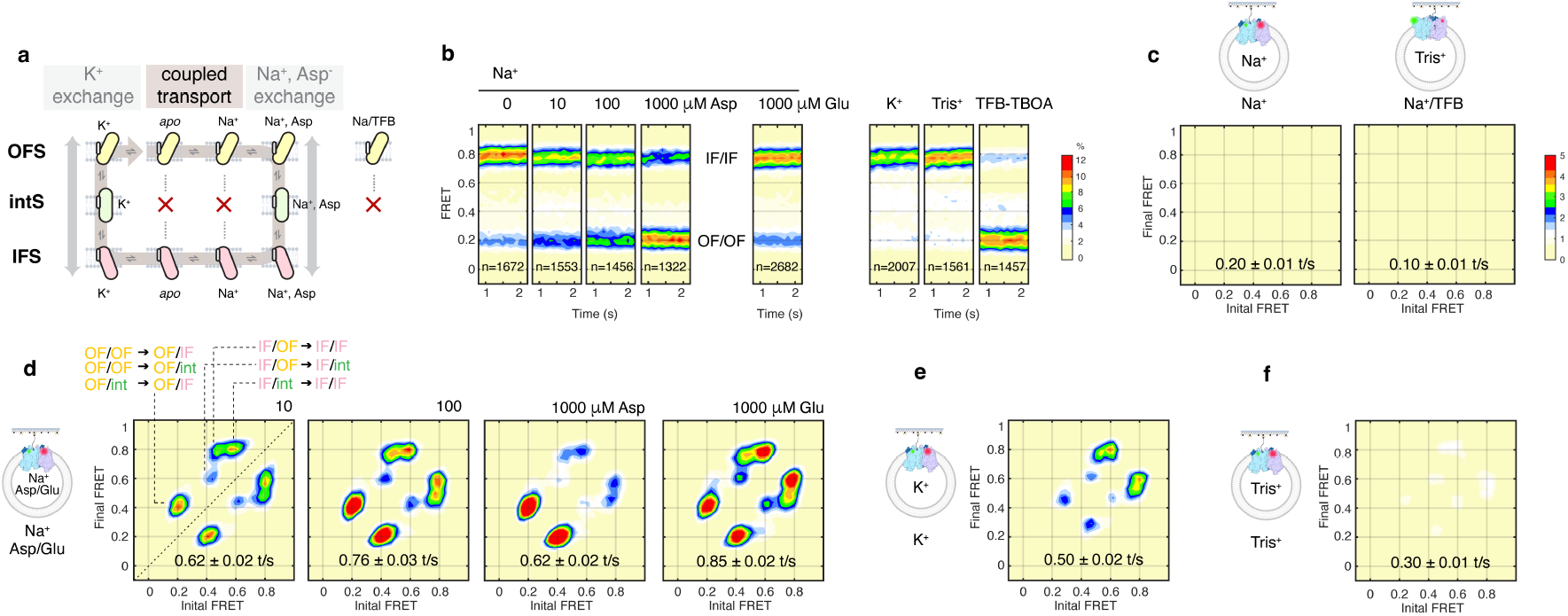
Ions and substrates control elevator dynamics. **a**, Transport cycle of EAAT1_fret_. Brown arrow indicates steps leading to concentrative substrate uptake in the presence of driving electrochemical gradients. Vertical arrows emphasize elevator modalities, during which Na^+^/Asp or K^+^ ions are exchanged reversibly across the membrane in the absence of ion gradients. Red crosses denote the absence of elevator transitions in the *apo* state, in the presence of Na^+^ ions alone, or in the presence of TFB-TBOA (TFB). **b**, FRET state distributions over time in the presence of (from left to right): 200 mM NaCl and the indicated concentrations of Asp or Glu; 200 mM KCl; alkali-free 200 mM TrisCl buffer; 200 mM NaCl and 400 µM TFB-TBOA. Total number of molecules analyzed (n) across three independent repeats shown on each panel. Scale bar shown on the right. **c-f**, Two-dimensional transition density plots of transitions between initial (abscissa) and final (ordinate) FRET states in the presence of 200 mM NaCl or 200 mM NaCl and 400 µM TFB-TBOA (**c**), 200 mM NaCl and the indicated concentrations of Asp or Glu (**d**), 200 mM KCl (**e**), and alkali-free 200 mM TrisCl (**f**). Scale bar (10^-3^ transitions per bin per second) shown on right in (**c**). Schematic experimental setups shown alongside: proteoliposomes (gray) immobilized on pegylated (black lines) imaging surface (blue) using a biotin (black dot)–neutravidin (light brown) bridge. Transition assignment indicated in leftmost panel of (**d**). Mean transition frequencies (number of transitions (t) per second) shown on plots. Data represent means ± SEM.

In the presence of symmetric 200 mM Na^+^ alone, EAAT1_fret_ predominantly stayed in the IF/IF state (FRET value of 0.8) and exhibited almost no dynamics, similar to when TFB-TBOA was present (**Fig. 2b, c**). When we added Asp or Glu on both sides of the membrane, we triggered Na^+^/Asp or Na^+^/Glu exchange and observed frequent elevator movements, manifesting as peaks in the transition density plots (**Fig. 2d**). Kinetic analysis showed that the substrates shortened the lifetimes of both IF/IF and OF/OF states (**ED Fig. 9s-t**), consistent with their catalyzing OF⇋IF transitions. These findings align with SLC1 structures, which show that helical hairpin 2 (HP2), which forms the substrate-binding gate, remains open when only Na^+^ is bound, thereby preventing elevator transitions. Subsequent substrate binding closes the gate and triggers the transitions ^34–38^. Strikingly, increasing Asp – less so for Glu – gradually shifted the transporter into the OF/OF state (0.2 FRET state, **Fig. 2b; ED Fig. 9q**) and decreased the number of observed transitions, which occurred mostly between OF/OF and OF/IF (0.4 FRET state, **Fig. 2d**). This is because Asp, unlike Glu, shortens the OF/OF lifetime significantly less than that of IF/IF (**ED Fig. 9s-t**). These results show that both substrates catalyze elevator dynamics in the presence of Na^+^, as expected, but also suggest that they can exhibit different transport kinetics.

With 200 mM K^+^ on both sides of the membrane, EAAT1_fret_ is inward-facing, populating the IF/IF state, and exhibits frequent transitions to all observable states (**Fig. 2b, e**). In contrast, elevator dynamics decrease in the absence of alkali ions (**Fig. 2f**), showing that K^+^ ions catalyze elevator transitions, as expected for K^+^ antiport. However, transitions are not completely eliminated without K^+^, and after perfusing alkali-free vesicles with external Na^+^ and TFB-TBOA, EAAT1_fret_ gradually shifts from mostly occupying the IF/IF state to the OF/OF state as the external blocker traps each protomer reaching OFS (**ED Fig. 7a-c**). These experiments show that EAAT1_fret_ protomers can move from IFS to OFS without antiported K^+^, although at slower rates. This is surprising because a tight coupling of the transport cycle to K^+^ antiport would require transitions to halt completely without K^+^. Thus, our results suggest that K^+^ is not strictly required for IFS-to-OFS transitions, and that the coupling of substrate uptake to K^+^ antiport is imperfect (**Fig. 2c,f**). Together, these experiments directly demonstrate that symported Na^+^ ions are transported only with the substrate, and that antiported K^+^ promotes the transporter’s return to the OFS.

### EAAT1 exhibits active and pausing modes

Examining individual smFRET traces recorded in symmetrical saturating Na^+^ and Glu, we observed striking heterogeneity, indicative of modal behavior (**Fig. 3a, ED Fig. 8-9**). Of 2682 recorded transporters, 17% displayed rapid transitions among all expected FRET states (**ED Fig. 9a**), which we interpret as both protomers undergoing elevator dynamics (*2-active*). Another 15% showed rapid transitions between FRET states of 0.2 and 0.4, or between 0.4, 0.6, and 0.8, consistent with one protomer being dynamic and the other paused in either the OFS or IFS, respectively (*1-active*). The mean transition rate of transporters in the *2-active* class was roughly twice that of the *1-active* class (**ED Fig. 9r**), suggesting similar dynamics across both groups. 13% of smFRET traces showed brief, infrequent transitions (fewer than 2 per second on average), suggesting largely quiescent protomers (2-*pausing*). Occasionally, we observed switching among 2-pausing, 1-active, and 2-active behaviors (**Fig. 3a** and **ED Fig. 8e**). Finally, 55% of traces remained static during the recording window, likely due to paused protomers. These were excluded from further analysis because their dwell lengths were determined by stochastic photobleaching and because they may contain damaged transporters or contaminants.

**Figure 3.**
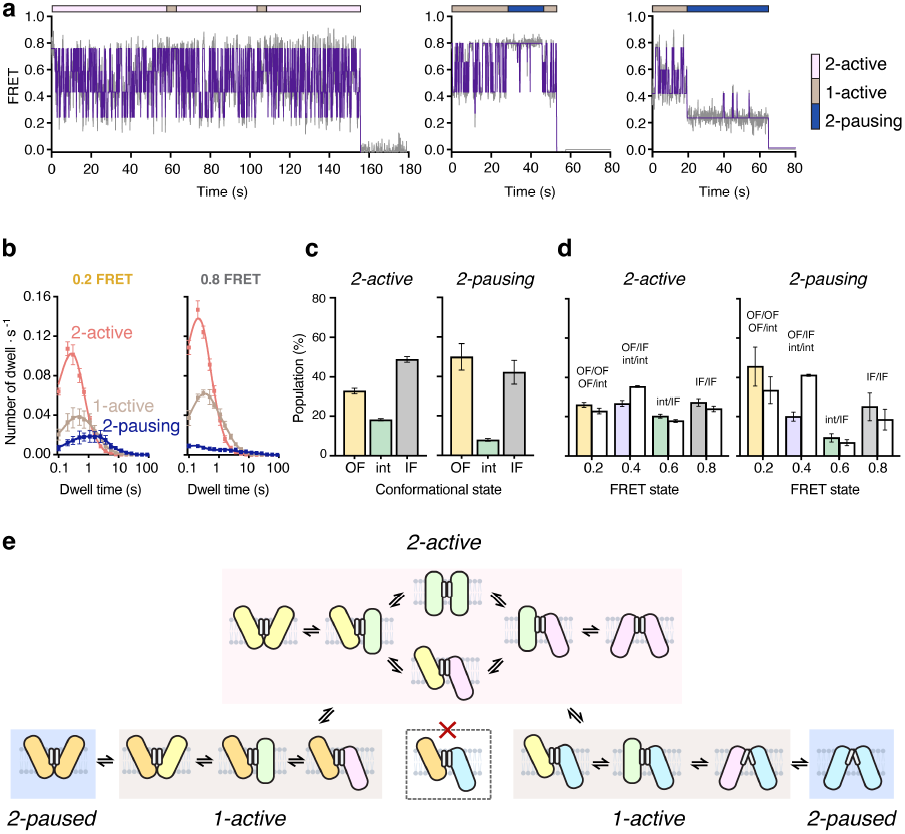
EAAT1_fret_ features multimodal dynamics. **a**, Representative smFRET traces of EAAT1_fret_ in symmetrical 200 mM NaCl and 1 mM Glu. Raw data, gray; idealization, purple. Horizontal bars indicate dynamic modes shown in color key on right. **b**, Dwell-time distributions for OF/OF (0.2 FRET, left) and IF/IF (0.8 FRET, right) states for 2-active (pink), 1-active (brown), and 2-pausing (dark blue) molecules. **c**, Deconvoluted protomer populations calculated for 2-active (left) and 2-pausing (right) molecules. OFS, yellow; intS, green; IFS, gray. **d**, FRET-state populations for 2-active (left) and 2-pausing (right) molecules measured experimentally (colors) and calculated from conformational populations in **c** assuming independent protomers (white). Data are means of three independent repeats ± SEM. **e**, Schematic representation of conformational changes in the two observed protomers in smFRET measurements.

To estimate the time spent in each conformation, we idealized individual smFRET traces and plotted dwell-time distributions for each kinetic class (**Fig. 3b**). Switching traces were classified by their dominant behavior. The combined dwell-time distributions were best fit with two or three components (**ED Fig. 9w**). Dwell times below 0.5 s correspond to the dynamic mode and mainly arise from 1-active or 2-active trimers (**ED Fig. 9w**). In contrast, longer dwell times mostly originate from 2-pausing trimers. Dwell times of the IF/int (0.6 FRET) state are close to 0.1 s (**ED Fig. 9w-z**), the temporal resolution of our recordings, indicating that intS is short-lived and that about half of the transitions into this state go undetected. Consistent with this, the transition density plots show ample direct transitions between OFS and IFS (0.4⇋0.8) (**Fig. 2d**), due to missed intS in the OFS⇋intS⇋IFS transitions (0.4⇋0.6⇋0.8).

Next, we calculated OFS, intS, and IFS populations from our dwell-time distributions (**Fig. 3c**). The intS population is smaller in the 2-pausing class than in the 2-active class, consistent with pausing occurring only in OFS or IFS, not in the transient intS. We then back-calculated the expected populations for the 0.2, 0.4, 0.6, and 0.8 FRET states, assuming that protomers behave independently. The FRET state distributions for the 2-active class were similar to those expected for independently functioning protomers (**Fig. 3d**). However, the FRET state distributions for 2-pausing molecules showed an underrepresentation of the 0.4 FRET state (**Fig. 3d**). These results suggest that pausing mostly occurs in symmetric OF/OF (0.2 FRET) and IF/IF (0.8 FRET) states, while avoiding the asymmetric OF/IF (0.4 FRET) state (**Fig. 3e**).

### Multimodal dynamics determine transport rates

We expect that modal elevator dynamics should produce modal transport activity, in which conformationally active molecules transport substrates quickly, while pausing molecules transport slowly. To test this, we combined smFRET-based dynamics measurements with single-transporter activity assays in proteoliposomes under non-equilibrium flux conditions. We first measured conformational dynamics under efflux and uptake conditions with varying ion and substrate gradients. First, vesicles loaded with 200 mM Na^+^ and 100 µM Asp were immobilized in smFRET chambers containing the same buffer (**Fig. 4a and ED Fig. 10a**). After initial recordings showing transporters mostly in the OF/OF (0.2 FRET) state (**Fig. 4b**), vesicles were perfused with 200 mM K^+^ to drive substrate efflux and mimic the pathological collapse of the Na^+^ and K^+^ gradients during ischemic stroke. About 60% of traces showed transporters shifting from OF/OF to other states (**Fig. 4c**); the remaining traces did not respond until photobleaching (**ED Fig. 10e**). 52% of the responding molecules exhibited ∼10 seconds of activity, sampling all FRET states, after which they mostly stopped, predominantly in the IF/IF (0.8 FRET) state. The activity duration was shorter when vesicles were loaded with 1 or 10 µM Asp and longer with 1000 µM Asp or Glu (**ED Fig. 10c-g**). The remaining 48% of molecules paused soon after reaching the IF/IF state. Mechanistically, these results suggest that, after K^+^ perfusion, EAAT1_fret_ protomers rapidly release bound Na^+^/Asp and transition from OFS through intS to IFS. Once in IFS, protomers bind Na^+^/Asp and either transition back to OFS in the next round of efflux or pause. When all Asp is transported out of the vesicles, the residual high internal Na^+^ inhibits further dynamics (**ED Fig. 10a**).

**Figure 4.**
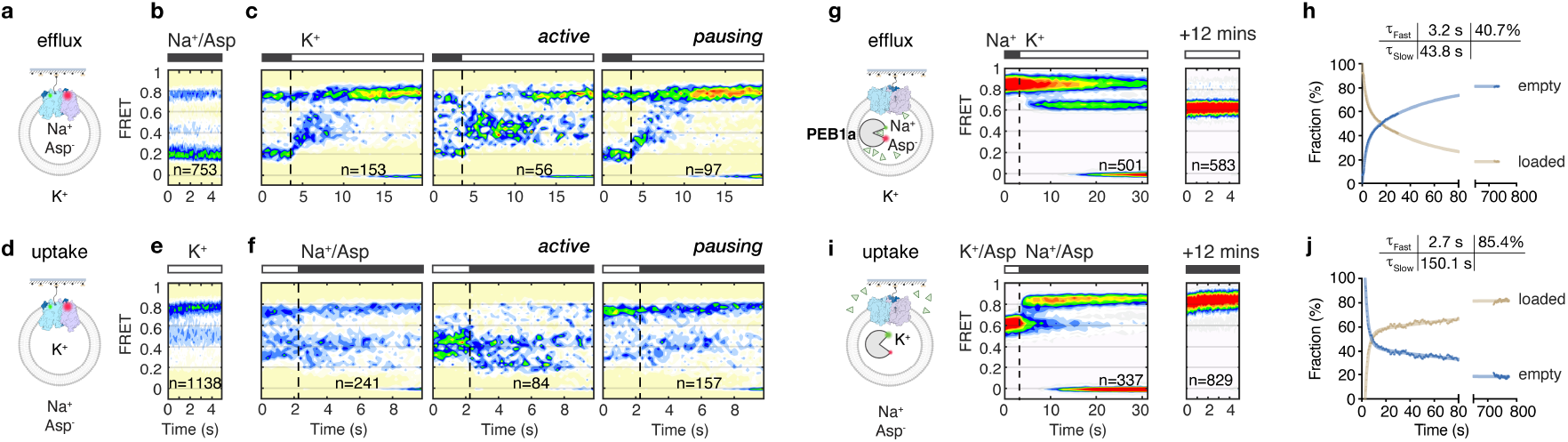
Transport dynamics reveal active and pausing modes. **a-f**, Single-molecule dynamics of LD555/LD655-labeled EAAT1_fret_ during Asp efflux (**a-c**) and uptake (**d-f**). Schematic experimental setups (**a**,**d**). FRET state distributions over time for EAAT1_fret_ prepared in 200 mM NaCl and 100 µM Asp (**b-c**) or 200 mM KCl (**e-f**). Initial recordings in symmetrical buffers permitting Na^+^/Asp (**b**) and K^+^ ion (**e**) exchange. Subsequent recordings during perfusion with 200 mM KCl (**c**) or 200 mM NaCl and 10 µM Asp (**f**) to initiate efflux and uptake, respectively. Dashed lines denote the onset of perfusion. Total molecules responded to perfusion (left); subset showing sustained activity (middle); subset that displays pausing features (right). **g-j**, Single-transporter assays of Asp efflux (**g**,**h**) and uptake (**i**,**j**) by EAAT1_fret_ proteoliposomes with the encapsulated LD555/LD655-labeled Asp sensor PEB1a. Schematic experimental setups (**g**,**j**) and FRET state population plots during real-time perfusion (middle) and after 12 minutes of equilibration (right). Bars above the plots indicate external buffers. Dashed lines denote the onset of perfusion. FRET of ∼0.8 corresponds to PEB1a bound to Asp (Asp-loaded vesicles); and FRET of 0.6 corresponds to unbound PEB1a (empty vesicles). Quantification of time-dependent changes in fractions of empty and loaded vesicles from fits to biexponential functions (**h**,**j**). Fast and slow time constants (*τ*) and corresponding populations shown in insets.

To measure conformational dynamics during substrate uptake, vesicles loaded with 200 mM K^+^ were immobilized and perfused with 200 mM Na^+^ and varying Asp concentrations (**Fig. 4d, ED Fig. 10b**). In K^+^ buffer, transporters mainly occupied the IF/IF (0.8 FRET) state, with smaller populations in other states (**Fig. 4e**). After Na^+^/Asp perfusion, 68% of traces showed transporters shifting toward OFS within 1 second. Of these, 41% remained active throughout the recording (**Fig. 4f, ED Fig. 10i**), consistent with continuous rounds of Na^+^/Asp transport. The remaining molecules entered a pausing mode after the initial FRET-state changes. Overall, we observed broad distributions of transition frequencies (**ED Fig. 10k**) under both efflux and uptake conditions, reflecting molecules with sustained activity and pausing (**ED Fig. 10l**,**m**).

We next measured substrate efflux and uptake using the single-transporter assay ^29^, in which proteoliposomes containing biotin-PEG_11_-maleimide-labeled EAAT1_fret_ were loaded with an Asp sensor – the Asp-binding protein PEB1a-Y198F, labeled with LD555 and LD655 dyes (**Fig. 4i**). During transporter reconstitution and sensor encapsulation, low protein concentrations were maintained, so that each vesicle contained no more than one transporter and sensor (see *Methods*). Labeled PEB1a-Y198F exhibits a 0.6 FRET state when unbound and a 0.8 FRET state when bound to Asp ^29^. To measure substrate efflux, we immobilized vesicles loaded with 200 mM NaCl and 100 µM Asp, corresponding to ∼25 Asp molecules in a typical ∼100 nm diameter vesicle (see *Methods*), and perfused them with 200 mM K^+^ (**Fig. 4g**). An initial FRET state of 0.8 indicated that PEB1a-Y198F was saturated with Asp. Given that PEB1a-Y198F has an affinity for Asp of ∼4 µM ^29^, the sensor is expected to shift to the 0.6 FRET state when the last Asp molecules are transported out of the vesicle. Approximately 41% of vesicles rapidly emptied their Asp, showing a transition from 0.8 to 0.6 FRET state with a time constant of ∼3.2 s (**Fig. 4h**). This indicates that individual protomers transported ∼2.6 Asp molecules per second, assuming each vesicle contains one EAAT1_fret_ trimer with three active protomers. The remaining vesicles took much longer to empty, with an estimated time constant of ∼43.8 s, consistent with them containing pausing transporters (**Fig. 4h**). To measure uptake, we perfused these emptied vesicles with 200 mM Na^+^ and 10 µM Asp (near the Michaelis constant for Asp transport) (**Fig. 4i**). Uptake of about three Asp molecules will saturate the sensor ^29^. About 85% of vesicles showed rapid uptake within 3 s, manifesting as a FRET change from 0.6 to 0.8 (**Fig. 4j**), showing that most EAAT1_fret_ trimers are able to mediate transport under appropriate ionic conditions. Together, conformational dynamics and transport assays strongly suggest that due to modal elevator dynamics, the transporters exhibit a mix of faster and slower transport kinetics.

### Structural substates correlate with activity

CryoEM of EAAT1_fret_ revealed substates within OFS and IFS, raising the question of whether they correlate with elevator dynamics. In the ^wrapped^OFS substate, a moiety modeled as cholesterol hemisuccinate (CHS) wedges between the transport domain and the TM4a helix of the scaffold (**Fig. 1d, 5a, and ED Fig. 6**). In contrast, ^free^OFS and other states of the transport cycle – intS and IFS substrates – lack CHS at this site, and their transport domains pack loosely against TM4a. The inter-domain interface is a hotspot for EAAT allosteric inhibitors that arrest elevator dynamics ^36,39^; therefore, we tested whether CHS slows elevator motions. We observed that increasing CHS from 0 to 33% reduced elevator dynamics (**Fig. 5b, j**) and favored the OF/OF (0.2 FRET) state (**Fig. 5c**) by lengthening dwell times of both its dynamic and pausing components (**Fig. 5d, f**), whereas it had little effect on IF/IF dwells (**Fig. 5e**). Notably, cryoEM imaging at ∼26% CHS showed ∼19% of protomers in the ^wrapped^OFS substate, suggesting that CHS binding is comparatively weak. These results suggest that CHS acts as a negative allosteric modulator of EAAT1_fret_ by selectively binding to ^wrapped^OFS.

**Figure 5.**
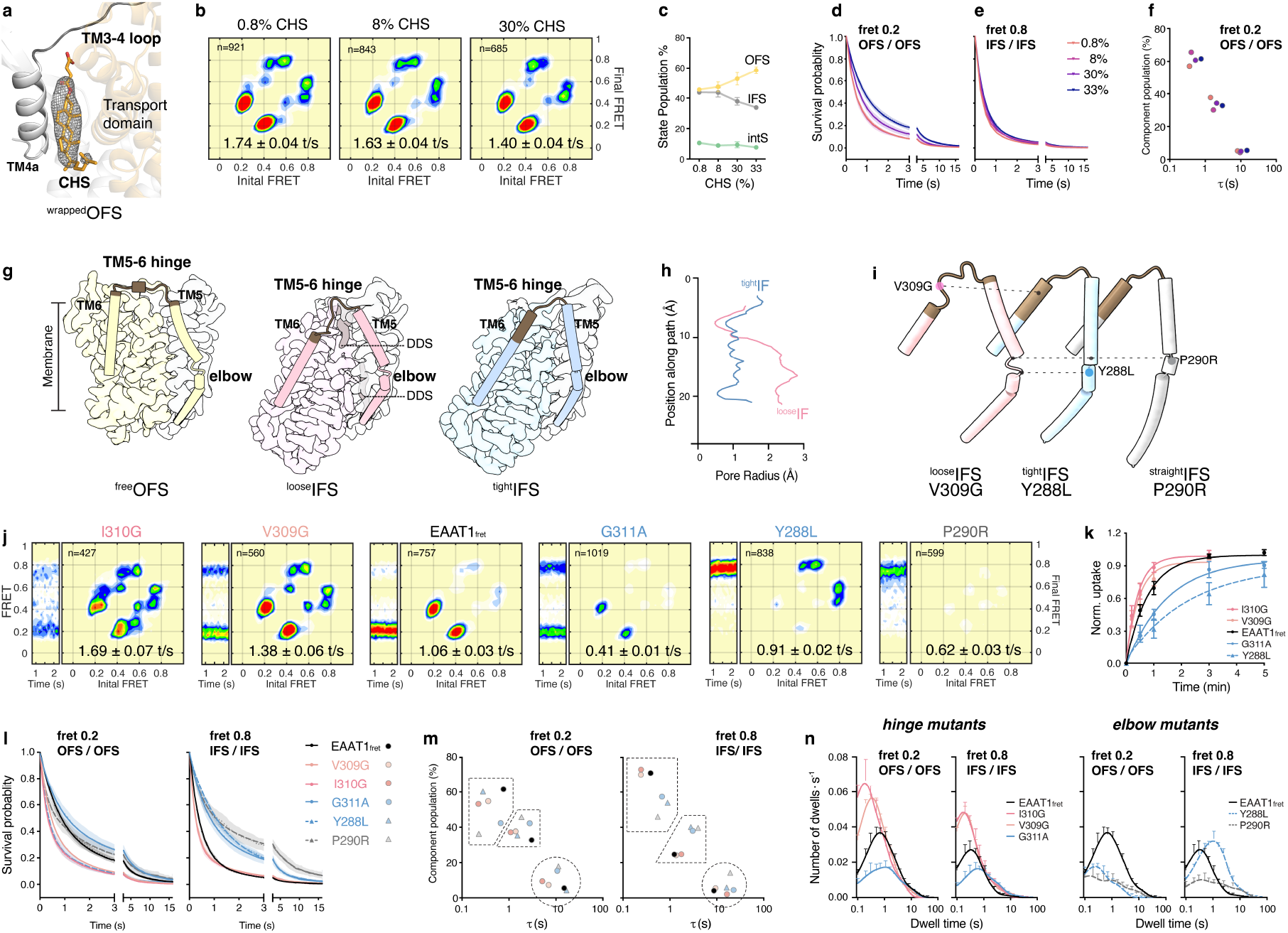
Membrane composition and mutations modulate transporter dynamics. **a**, Close-up of the TM3-4 hinge in the ^wrapped^OFS of EAAT1_fret_, showing cryoEM density for CHS. Scaffold TM4a, grey; transport domain, orange. **b**, Transition density plots for proteoliposomes containing 0.8, 8, or 30% CHS; data for 33% CHS are in **j**. Molecule numbers (*n*) and mean transition frequencies indicated. **c**, CHS-dependent deconvoluted populations of protomers in OFS, intS, and IFS. **d-e**, Survival probability for OF/OF (0.2 FRET) (**d**) and IF/IF (0.8 FRET) (**e**) states fitted to triexponential functions. **f**, Fitted time constants (abscissa) and fractions (ordinate) of the three kinetic components for each curve in (**d**). Solid circles colored according to CHS content in (**d**). **g**, CryoEM maps of ^free^OFS, ^loose^IFS, and ^tight^IFS protomers. TM5 (scaffold domain) and TM6 (transport domain) shown as cartoons; hinges highlighted in brown. **h**, Pore radii at the domain interface of ^loose^IFS (pink) and ^tight^IFS (blue) calculated with HOLE. **i**, TM5-TM6 structures of V309G, Y288L, and P290R mutants. **j**, FRET populations over time (left) and transition density plots (right) for constructs with decreasing activity (left to right): I310G, V309G, EAAT1_fret_, G311A, Y288L, and P290R. Molecule numbers (*n*) and mean transition frequencies indicated. **k**, L-[^3^H]aspartate uptake over time in HEK293 cells expressing EAAT1_fret_ and its mutants. Data represent at least 5 replicates for each transporter from 2-3 independent transfections. **l**, Survival probability for OF/OF (0.2 FRET, left) and IF/IF (0.8 FRET, right) states fitted to triexponential functions. **m**, Time constants (abscissa) and fractions (ordinate) corresponding to fitted values in (**l**). Solid circles (hinge mutants) and triangles (elbow mutants) colored as in (**k**). Dotted lines serve to visually separate the three kinetic components. **n**, Dwell-time distributions of OF/OF and IF/IF states for hinge mutants (solid lines) and elbow mutants (dashed lines). Data are means of three independent replicates ± SEM. Number of transporters analyzed indicated in **j**.

In detergent-solubilized IFS, we noted three differences between the ^loose^IFS and ^tight^IFS substates. First, the transport and scaffold domains are tightly packed in ^tight^IFS but loosely packed in ^loose^IFS (**Fig. 5h**), with the hydrocarbon moieties of the detergent sucrose monododecanoate (DDS) inserted at the interface (**Fig. 5g**). ^loose^IFS resembles the “unlocked” state of the archaeal homolog Glt_Ph_, which exhibits rapid elevator dynamics ^27^. Second, the TM5-6 hinge (residues 307-312) that connects the scaffold to the transport domain is an extended loop in ^loose^IFS, as in OFS and intS, but adopts a helical conformation that extends TM6 in ^tight^IFS (**Fig. 5g**). ^loose^IFS itself comprises an ensemble of smaller substates, differing in hinge details and the degree of transport domain tilt (**ED Fig. 6**). Third, in the scaffold domain, a flexible “elbow” linking a central trimerization region with a more flexible peripheral helical bundle containing the C-terminal part of TM5 is sharply bent in ^loose^IFS, OFS, and intS, but straighter in ^tight^IFS (**Fig. 5g, ED Fig. 6b**). Strikingly, a P290R mutation linked to human episodic ataxia type 6 ^14^ is located at this elbow, suggesting it is functionally critical.

Hypothesizing that these differences might make ^loose^IFS more dynamic than ^tight^IFS, we introduced mutations V309G and I310G to increase hinge flexibility, G311A to increase helicity, and Y288L and P290R to influence helix bending at the TM5 elbow (**Fig. 5i-n**). As anticipated, V309G showed only ^loose^IFS in cryoEM (**Fig. 5i, ED Fig. 6**), whereas both ^loose^IFS (∼40%) and ^tight^IFS (∼60%) were observed for WT EAAT1_fret_. Both V309G and I310G increased smFRET dynamics and EAAT1_fret_-mediated Asp uptake in cells, while G311A decreased uptake (**Fig. 5j, k**). In smFRET traces, V309G and I310G accelerated both the fast and slow components in the OF/OF state and slightly shortened the fast component in the IF/IF state (**Fig. 5l, m**), resulting in left-shifted dwell-time distributions, increased abundance of dwells shorter than 0.5 s, characteristic of the dynamic mode (**Fig. 5n**), and more elevator transitions (**Fig. 5j**). In contrast, G311A slowed the fast and slow components in the IF/IF state and increased the fraction of the slow components in both the IF/IF and OF/OF states. Elbow mutations, Y288L and P290R, reduced dynamics and Asp uptake (**Fig. 5j, k**), and cryoEM showed that Y288L was present only in ^tight^IFS (**Fig. 5i, ED Fig. 6**). Like G311A, they lengthened the fast and slow kinetic components of the IF/IF state and increased the fraction of slow components (**Fig. 5l-m**), resulting in a rightward shift in the IF/IF dwell time distributions (**Fig. 5n**). Unexpectedly, Y288L shortened OF/OF state dwell times (**Fig. 5l, m**) but also significantly reduced transitions into the OFS (**Fig. 5j, n**), with most transitions occurring between intS and IFS (**Fig. 5j**); these transitions likely do not contribute to substrate transport since they do not complete the transport cycle.

Together, these results show that factors such as lipid binding and mutations that favor ^wrapped^OFS or ^tight^IFS substates slow elevator dynamics, whereas those that favor ^free^OFS or ^loose^IFS increase dynamics. Rigidification of the inter-domain hinges, straightening of the scaffold elbow, and more extensive domain packing correlate with fewer elevator transitions and longer pauses.

### Molecular basis of episodic ataxia type 6

P290R is associated with the most severe episodic ataxia type 6 phenotype reported to date. Impaired surface expression, loss of substrate transport, and dramatically increased Cl^-^ conductance constitute the pathophysiology of P290R ^12–14,40–42^, but the molecular mechanisms underlying these changes remain unclear. We therefore leveraged our integrated platform to examine the structure and dynamics of P290R in detail. In smFRET experiments, the mutant was mostly trapped in the IF/IF state and showed few transitions (**Fig. 5j**), consistent with the loss of transport. Similarly, cryoEM structures of P290R in detergent or nanodiscs revealed only protomers in the IFS. In detergent, we observed an asymmetric trimer, with each protomer in an unusual IFS conformation (**ED Fig. 11**): two resembling the ^tight^IFS, and the third showing an even straighter TM5 (^straight^IFS) (**Fig. 5i**). In nanodiscs, a symmetric trimer features protomers similar to the ^straight^IFS but with their transport domains shifted slightly outward (^straight^IFS*). This straighter elbow was expected, as the arginine residue replaces the helix-disrupting proline. Structural changes at the elbow propagate through the rest of the scaffold domain, which hugs the transport domain and increases the interface from ∼1700 Å^2^ in ^tight^IFS to ∼1950 Å^2^ in ^straight^IFS*. In contrast, the trimerization interface is distorted, with three well-packed F270 residues on the symmetry axis replaced by L267 residues (**ED Fig. 11a-c**), which may explain the reduced stability and cell expression of this mutant.

The Cl^−^-conducting pathway in EAAT1 opens between the transport and scaffold domains in the transient chloride-conducting state, which is structurally similar to the IFS ^43^. In the IFS substates of EAAT1_fret_, the entire pathway is tightly constricted. In contrast, in the P290R ^straight^IFS*, the pathway is more open (**ED Fig. 11f**). Although the extracellular gate residues, L296 and M89, still constrict the pathway, the intracellular constriction, formed by M286 and Q445, is widened (**ED Fig. 11e**,**f**). The proximal guanidinium group of R290 creates a local electropositive environment at the cytoplasmic entrance, potentially increasing attraction for Cl^−^ (**ED Fig. 11h**).

Interestingly, we did not observe similarly dramatic structural consequences of the equivalent mutation, P259R, in EAAT3, which produces a pathological phenotype similar to that in EAAT1 in cell-based studies ^42^. In detergent, WT EAAT3 predominantly adopts the ^open^IFS state, with two Na^+^-binding sites occupied, an empty substrate-binding site, and an open HP2 gate, reflecting the low affinity of this conformation for Asp ^34^. The P259R mutant produced three IFS conformations: an Asp-free ^open^IFS, similar to WT, and two Asp-bound classes with closed HP2 (^closed^IFS and ^closed^IFS*; **ED Fig. 11k-l**), consistent with the mutation increasing substrate affinity, as in EAAT1 ^12^. In neither class was the scaffold domain reordered, as in EAAT1_fret_ P290R (**ED Fig. 11i-j**). Similarly, the equivalent mutation in an archaeal homolog was insufficient to restructure the scaffold ^44^. These results suggest that the structural rearrangements of the scaffold domain are a concerted event, governed by the energetics of the entire domain rather than by the elbow alone.

Overall, our findings suggest that the ataxia type 6 phenotype in the EAAT1 P290R mutant arises because the mutation favors a distinct scaffold domain conformation with a straightened elbow, an increased interaction surface with the transport domain in IFS, disrupted trimerization interactions, and a greater propensity to open the Cl^-^ permeation pathway.

## Discussion

EAATs have emerged as potential pharmacological targets for a wide range of neurological disorders, including neuropathic pain, epilepsy, neurodegenerative disorders, and addiction. In all cases, the therapeutic goal is to enhance Glu reuptake into glial cells to dampen excitatory neurotransmission or prevent excitotoxicity. Although promising, the development of successful small-molecule activators ^20,24,25^ has been hindered by a limited understanding of their mechanism of action. We previously used smFRET-based dynamics ^26,27,30^ and single-transporter activity assays ^29,45^ to identify which transport steps of an archaeal EAAT homolog, Glt_Ph_, are rate-limiting and how mutations affect them, thereby proposing a mechanism by which activators might act. However, this level of understanding remained beyond reach for human transporters. The development of a thermally stabilized human EAAT1 variant ^31^ provided the breakthrough that enabled us to establish an experimental platform to study the conformational dynamics underlying transport activity by integrating smFRET, single-molecule transport assays, and cryoEM.

Our combined smFRET and cryoEM experiments allowed us to assign most observed FRET states, enabling the study of conformational transitions between them with sub-second temporal resolution. These measurements revealed the dynamic basis for coupled transport of substrates and co- and counter-transported ions. EAAT1 elevator motions are rapid when the transporter is loaded with substrate and co-transported Na^+^, but nearly halted with Na^+^ alone, thereby ensuring substrate and Na^+^ symport and preventing Na^+^ leaks. Similarly, K^+^ stimulates the dynamics of the empty *apo* transporter, ensuring K^+^ counter-transport during the return step of the transport cycle. The high resolution of our measurements also revealed coupling imperfections: *apo* transporters can still move between the IFS and OFS, albeit at slower rates. Similar observations were made for the multidrug EmrE transporter ^46,47^, in which ion abundance affects the stoichiometry of coupling. We furthermore observed the importance of energetic balance between the OFS and IFS. Mutations that introduce an imbalance, such as IFS-stabilizing Y288L and P290R, diminish transport activity by preventing the transporter from reaching the OFS. Surprisingly, Asp, but not Glu, favors OFS, leading to depressed elevator dynamics at high Asp concentrations.

Strikingly, individual smFRET traces of EAAT1_fret_ revealed that sustained high activity is interspersed with periods of low activity, marked by long pauses and infrequent, brief transitions. This behavior was observed under both equilibrium conditions and in the presence of ion and substrate gradients, leading to substrate uptake or efflux from lipid vesicles. Consistent with this, single-molecule assays monitoring the activity of individual transporters also indicated a mixture of transporters operating at rates that differed by an order of magnitude. Elevated membrane CHS, which favors ^wrapped^OFS, and mutations that favor ^tight^IFS, with a rigidified TM5-6 hinge and a straightened scaffold elbow, slow elevator dynamics during active periods and promote pausing. Mutations that increase hinge flexibility and favor ^loose^IFS increase elevator dynamics but do not diminish pausing. Thus, we conclude that entry into ^wrapped^OFS and ^tight^IFS slows overall dynamics, while entry into ^free^OFS and ^loose^IFS increases dynamics. Yet the structural basis of pausing remains incompletely understood, as with modal activity observed in channels, pumps, and enzymes ^48–50^. Nevertheless, we attribute pausing to specific structural states that act as kinetic traps (**Fig. 6**). We therefore propose that allosteric activators of EAATs can be developed that either accelerate transport dynamics during active periods or reduce the probability of pausing, for example, by preferentially binding to active conformations.

**Figure 6.**
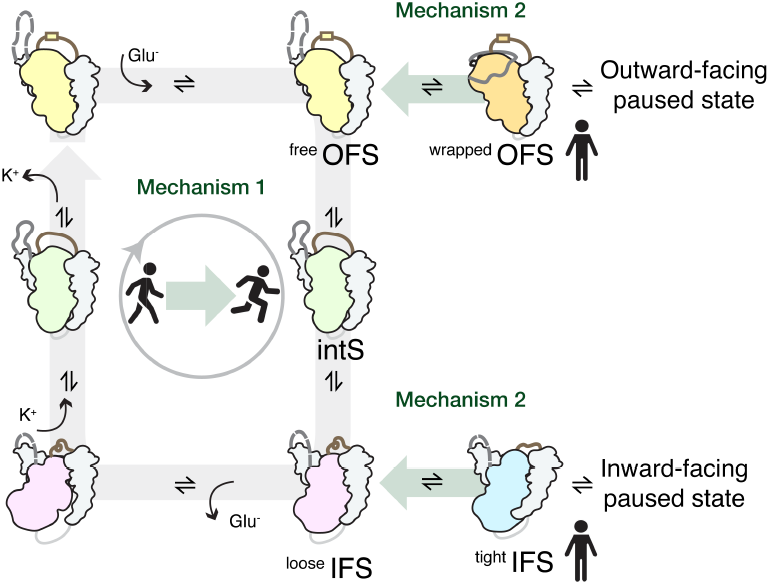
Paused states slow EAAT1_fret_ transport. Schematic representation of the EAAT1_fret_ transport cycle with ^wrapped^OFS and ^tight^IFS shown as off-cycle intermediates that might lead to paused states. Positive allosteric modulators may act by increasing the rate of transport in the active cycle or by disfavoring states that lead to pausing.

When active, protomers in EAAT1_fret_ trimers move independently. However, when pausing, the two observed protomers mostly pause in either OFS or IFS, avoiding asymmetric OF/IF states. This was a surprising result, suggesting allosteric communication between protomers in the trimer. By contrast, the archaeal homolog Glt_Ph_, which also shows modal dynamics, does not exhibit subunit coupling – its protomers move independently, even as they transition between fast and slow dynamic modes ^26,27,51,52^. In Glt_Ph_, this is likely due to a highly rigid scaffold that limits allosteric communication between subunits. Allosteric communication via the membrane has also been ruled out for this transporter ^53,54^. In EAAT1, however, the scaffold domain is more flexible, and we have identified OFS and IFS substates that differ in scaffold structure and the connecting hinges; mutations that affect transport dynamics also affect these regions. P290R is particularly consequential, resulting in a drastic restructuring of the scaffold that affects both its interactions with the transport domain and its trimerization.

Using our platform, we examined the mechanistic origin of a human neurological disorder. EAATs function as substrate-gated Cl^-^ channels ^2,55^ in addition to mediating ion-coupled transport, a feature conserved from prokaryotes to mammals ^56^. Pathological increases in this Cl^-^ conductance underlie episodic ataxia caused by the P290R mutation in human EAAT1 ^12,13^. A Cl^-^ channel opens between the scaffold and transport domains in a transient Cl^-^-conducting state structurally similar to IFS ^43^. Our smFRET data show that P290R dramatically extends the IFS lifetime, perhaps allowing more excursions into the Cl^-^-conducting state, which would explain the reported increase in channel open probability ^13,42^.

In summary, we have established an experimental platform to visualize the structure and dynamics of the human glial transporter EAAT1 at single-molecule resolution. Our results have revealed the basis of ion coupling, the energetic and kinetic features of transport, and how these features can be altered by environmental factors and mutations. This platform sets the stage for future studies of transporter regulation and for identifying small-molecule modulators.

## Methods

### Protein expression and purification

All protein constructs were cloned into the modified pCDNA 3.1 vector (*GenScript*) with an N-terminal Strep-II tag followed by superfolder green fluorescent protein (GFP) and the PreScission protease site. The EAAT1_fret_ construct for FRET measurements was based on the thermally stabilized human _coco_EAAT1^31^, with N422C and other mutations introduced by site-directed mutagenesis and sequence-verified (*GenScript*). EAAT1_fret_ and its V309G, I310I, G311A, Y288L, and P290R mutants were purified as previously described^31^. In brief, EAAT1_fret_ constructs were expressed in suspension FreeStyle™ 293-F cells. Cells were cultured in Freestyle 293 medium (*Gibco*) to a density of approximately 1.2 million cells/mL, then transfected with 2 mg of plasmid using 25k linear polyethylenimine (PEI, *Polysciences*) at a 1:3 plasmid-to-PEI weight ratio. After 20 hours, transfected cells were supplemented with 2.2 mM sodium valproate (*Sigma-Aldrich*) to boost expression. Approximately 50 hours post-transfection, cells were collected by centrifugation at 4,000 g for 20 min at 16 °C. Cell pellets were resuspended in lysis buffer containing 50 mM HEPES/Tris, pH 7.4, 1 mM L-aspartate, 1 mM EDTA, 1 mM tris(2-carboxyethyl) phosphine (TCEP), 1 mM phenylmethylsulfonyl fluoride (PMSF), and a 1:200 dilution of protease inhibitor cocktail (*MilliporeSigma*). Resuspended cells were disrupted using an EmulsiFlex-C3 cell homogenizer (*Avestin*). Cell debris was removed by centrifugation at 4,000 g for 30 min at 4 °C, and membrane pellets were collected by ultracentrifugation at 186,000 g for 1 hour at 4 °C. Membrane pellets were homogenized in lysis buffer supplemented with 200 mM NaCl and 20% (v/v) glycerol (solubilization buffer) and incubated with 2% sucrose monododecanoate (DDS; *Anatrace*) and 0.4% cholesteryl hemisuccinate (CHS; *Anatrace*) on a rotator at 4 °C for 2 hours. Insoluble material was removed by ultracentrifugation at 186,000 g for 1 hour at 4 °C, and the supernatant was diluted 5-fold with dilution buffer (solubilization buffer free of glycerol, DDS, and CHS) and incubated with Strep-Tactin Sepharose resin (*GE Healthcare)* for 1 hour at 4 °C. The resin was washed with seven column volumes of wash buffer containing 50 mM HEPES/Tris, pH 7.4, 200 mM NaCl, 0.048% DDS, 0.012% CHS, 1 mM TCEP, 2.5% glycerol, and 1 mM L-aspartate. Protein was eluted with five column volumes of the wash buffer supplemented with 5 mM D-desthiobiotin (elution buffer). The Strep-II and GFP tags were cleaved by incubating the protein with homemade PreScission protease at a 40:1 protein-to-protease ratio overnight at 4 °C. The protein was further purified by size exclusion chromatography (SEC) using a Superose 6 Increase 10/300 column (*GE Healthcare*) pre-equilibrated with the desired buffer.

The EAAT3 mutant P259R was prepared in the de-glycosylated (N178T, N195T) construct as previously described ^32,34,39^. The procedures were similar to those as described above for EAAT1_fret_, except: a) 1 L suspension FreeStyle™ 293-F cells at a density of 2.5 million cells/mL were transiently transfected with 3 mg DNA and 9 mg PEI 25 K (Polysciences) and diluted to ∼1.2 million cells/ml after 6 hours; b) instead of DDS, the detergent Dodecyl-β-D-maltopyranoside (DDM, Anatrace) was used for solubilization, 0.06% glycodiosgenin (GDN, Anatrace) was used in wash and elution, and 0.01% GDN was used in SEC.

### Cryo-EM sample preparation

For the detergent samples, 3.5 μl of purified protein at 3-6 mg/ml was applied to glow-discharged Quantifoil R1.2/1.3 holey carbon-coated 300-mesh gold grids (*Quantifoil*). The grids were blotted for 3 s and plunge-frozen into liquid ethane using an FEI Mark IV Vitrobot (*Thermo Fisher Scientific*) at 4 °C and 100% humidity.

For the TFB-TBOA sample, EAAT1_fret_ at 6mg/ml (purified with 200 mM NaCl and 1 mM L-aspartate as described above) was diluted 200-fold into buffer containing 200 mM NaCl and 0.4 mM TFB-TBOA, 50 mM HEPES/Tris, pH 7.4, 0.048% DDS, 0.012% CHS, 1 mM TCEP, 2.5% glycerol. After incubating on ice for 30 minutes, the sample was concentrated to 3 mg/ml and plunge-frozen into liquid ethane as described above.

For nanodisc samples, Membrane Scaffold Protein (MSP2N2) was expressed and purified as previously described^57^. A mixture of 1-palmitoyl-2-oleoyl-glycero-3-phosphocholine (POPC), 1-palmitoyl-2-oleoyl-sn-glycero-3-phospho-L-serine (POPS), and cholesteryl hemisuccinate (CHS) (*Avanti Polar Lipids*) at a 3:1:0.4 (w:w) ratio was dried, hydrated in 50 mM HEPES/Tris, pH 7.4, supplemented with 200 mM NaCl and 1 mM L-aspartate for substrate-bound datasets, or 200 mM KCl for the *apo* dataset, and solubilized in 80-90 mM DDS with 8-9 mM CHS at a final lipid concentration of 20-30 mM (∼26% CHS, w/w lipids). EAAT1_fret_ constructs with Strep-II and GFP tags were mixed with MSP2N2 and solubilized lipids at a molar ratio of 1:1.5:62, aiming for a final lipid concentration greater than 5 mM. This mixture was incubated at 22 °C for 45 minutes, then 400 mg BioBeads SM-2 (*BioRad*) per mL mixture (w/v) was added for one hour at 22 °C with rotation, followed by a second round overnight at 4 °C on a rotator, and a last round of 200 mg BioBeads for one hour at 4 °C. Empty nanodiscs were cleared from the resulting MSP2N2/protein preparations by affinity chromatography using the same protocol as above. The Strep-II and GFP tags of the concentrated eluted product were cleaved by incubating with homemade PreScission protease at a 40:1 protein-to-protease ratio at 4 °C for three hours, followed by ultracentrifugation at 100,000 x g for 30 minutes. The EAAT1_fret_ proteins in MSP2N2 nanodiscs were applied to a Superose 6 Increase 10/300 column in 50 mM HEPES, pH 7.4, 1 mM TCEP, 2.5% glycerol, and supplemented with 200 mM NaCl and 1 mM L-aspartate for substrate-bound datasets, or 200 mM KCl for the *apo* dataset. The peak fractions were concentrated to 2-4 mg/mL. To improve particle distribution, fluorinated Fos-Choline-8 (*Anatrace*) was added to a final concentration of 1.5 mM. UltrAuFoil R1.2/1.3 300-mesh gold grids (*Quantifoil*) were glow-discharged at 25 mA for 80 s. 3.5 µL of protein was applied to each grid, incubated for 40 s under 100% humidity at 16 °C, blotted for 3 s, and plunge frozen in liquid ethane using a Vitrobot Mark IV.

### Cryo-EM data acquisition and image processing

Grids were screened on the Glacios microscope with the Falcon 4i camera at the WCM Cryo-EM facility. High-resolution imaging using Leginon software ^58^ was performed on the Titan Krios microscope with a Gatan K3 imaging system at the Janelia cryoEM facility (V309G dataset), the Columbia University Medical Center Cryo-EM core (Y288L dataset), and New York University Langone’s Cryo-EM laboratory (all other datasets). Data were collected at nominal magnifications of 105,000X or 85,000X. Microscopes were equipped with either a 15 eV or a 20 eV energy filter. Specific details for each data collection, as well as individual maps and models, are provided in **ED Table 1**. Visual summaries of processing workflows and EM validation are provided in **ED Figs 3-5**.

In a typical processing workflow, raw movies were imported into cryoSPARC 4.0-4.4 and subjected to Patch Motion Correction and Patch CTF Estimation, with the movies binned 2-fold ^59^. Particles were picked with the blob or template picker in cryoSPARC or with Warp ^60^ and extracted with a box size of 300 to 416 pixels, and then binned to 128 pixels. The particles were cleaned by 2D classification and used in *ab initio* reconstruction and nonuniform refinement ^61^ with C1 symmetry to generate a good template. 6-8 decoy volumes were obtained by running one iteration of *ab initio* reconstruction. Particles retained after 2D selection, which removed obvious non-protein junk, were cleaned by heterogeneous refinement using one good template and 4-6 decoy noise volumes. When greater than 95% of particles converged into the protein volume, the remaining particles were re-extracted to full box size and subjected to nonuniform refinement with C1 symmetry for the trimer map and with C3 symmetry to generate the consensus volume. Symmetry expansion and focused 3D classification were then performed using a single protomer mask. Each class underwent local refinement using pose/shift Gaussian priors during alignment (standard deviations of 3º, 2 Å).

Some datasets benefited from particle polishing. The particle coordinates were converted to Relion format using PyEM ^62^. Particles were re-extracted in Relion from motion-corrected movies binned 2x using MotionCor ^63^. These particles were re-imported into cryoSPARC, refined using nonuniform refinement (C1), converted again to Relion format with PyEM, and subjected to Bayesian Polishing. The polished particles were then subjected to heterogeneous refinement, followed by nonuniform refinement to generate the consensus volume.

### Model building and refinement

Unsharpened, cryoSPARC-sharpened, and DeepEMhancer-sharpened maps ^64^ were used for model building. EAAT1_fret_ structural models (**ED Table 1**) were fitted into EM density maps using ChimeraX ^65^. The models were manually adjusted in COOT ^67^ and ISOLDE ^68^ and subjected to rounds of real-space refinement in Phenix ^69^ using unsharpened density and default parameters. Structural model validation was performed in Phenix (**ED Table 1**). All structural figures were prepared using ChimeraX ^65^ or PyMOL ^70^. Most structural biology applications used in this project were compiled and configured by SBGrid ^71^.

### EAAT1_fret_ labeling for single-molecule FRET assays

For the dynamics assay, EAAT1_fret_ and mutant constructs were cleaved with PreScission protease to remove Strep-II and GFP tags, then SEC-purified in 50 mM HEPES/Tris, pH 7.4, 200 mM NaCl, 0.048% DDS, 0.012% CHS, 2.5% glycerol, and 1 mM L-aspartate. Proteins were labeled with LD555-MAL and LD655-MAL dyes (*Lumidyne Technologies*) and biotin-PEG_11_-maleimide (*EZ-link, Thermo Fisher Scientific*) at a molar ratio of 1:2.7:1.3:0.75. Excess reagents were removed using a 100-kDa centrifugal concentrator, and the labeled EAAT1_fret_ transporters were further purified by SEC on a Superose 6 Increase 10/300 column (*GE Healthcare*). Labeling efficiency was estimated by SEC with absorbance detection at 280, 550, and 650 nm for protein, LD555, and LD655, respectively.

For the transport assay, EAAT1_fret_ was labeled with biotin-PEG_11_-maleimide in the presence of N-ethylmaleimide (NEM) at a molar ratio of 1:2:0.5 (protein:biotin:NEM), as previously described ^29^.

### Labeled EAAT1_fret_ reconstitution into liposomes and ccPEB1a-Y198F encapsulation

Chloroform solutions of POPC and POPS at a 3:1 (w/w) ratio were mixed with or without CHS (10% of total lipids) to achieve 0% and 10%CHS liposomes. The lipid mixture was dried under N_2_ on a rotary evaporator and then left under vacuum overnight. The lipids were hydrated with 50 mM HEPES/Tris, pH 7.4, supplemented with the desired salts and amino acids, and subjected to 10 freeze/thaw cycles at a final concentration of 4 mg/ml. The suspensions were extruded through polycarbonate filters with 100-nm pores (*Whatman*) 11 times. The resulting liposomes were destabilized with 1% DDS, and the labeled EAAT1_fret_ proteins were added at a 1:100 (w:w) protein-to-lipid ratio (PLR) and incubated for 50 min at room temperature. At this destabilizing step, the reconstitution reaction with 0% or 10% CHS/lipid mixtures was supplemented with 2 mg of solubilized CHS to achieve 33% and 40% CHS in the reconstituted proteoliposomes, respectively. Detergents were removed by six rounds of incubation with 100 mg/ml pre-washed Bio-Beads SM-2 (*Bio-Rad Laboratories*): 2 hr at room temperature, 2 hr at 4 °C, overnight at 4 °C, and three rounds of 2 hr at 4 °C with gentle agitation. The proteoliposomes containing labeled EAAT1_fret_ were aliquoted into small quantities for immediate use or flash-frozen in liquid nitrogen for future use.

A glutamate/aspartate sensor, ccPEB1a-Y198F, was purified by His-tag affinity chromatography and labeled with LD555-MAL and LD655-MAL dyes as described previously^29^. The sensor was added to the EAAT1_fret_-containing proteoliposomes at a final concentration of 0.6 μM and encapsulated by three freeze/thaw cycles. Unencapsulated ccPEB1a-Y198F was removed by PD MiniTrap G-25 columns, and the proteoliposome sample was extruded 11 times through 100-nm-pore polycarbonate filter membranes (*Whatman*) before use.

### Determining lipid composition by NMR

The final CHS content in proteoliposomes was determined by NMR spectroscopy (**ED Fig 13**). 100-300 μL of proteoliposome samples were lyophilized and then dissolved in 500 μL of deuterated chloroform (99.8 atom % D, *Sigma Aldrich*). The suspension was vortexed vigorously and centrifuged at 22 kg at 4 °C. 200 μL of the supernatant was transferred to a 3-mm NMR tube for measurement. ^1^H-NMR spectra were collected on a Bruker Advance III HD 500 MHz spectrometer equipped with a TCI ^1^H-^19^F/^13^C/^15^N triple-resonance cryogenic probe (*Bruker*). The standard zg30 pulse sequence was used, and D1 was set to 1 s. The spectra were processed using Topspin 3.6.3 (*Bruker*). After baseline correction, the peaks at 3.32 ppm, corresponding to the -N(CH_3_)_3_ group of POPC, and 0.7 ppm, corresponding to the 18-CH_3_ group of CHS, were selected and integrated (**ED Fig 13a-b**). The ratio of the integrated areas was used to calculate the lipid-to-CHS ratio, assuming a 3:1 POPC:POPS ratio (**ED Fig 13c**). The measured CHS content of nominally 0, 10, 33, and 40% proteoliposomes was about 0.8, 8, 30, and 33%, respectively. The non-zero CHS content in the nominally 0% CHS proteoliposomes is likely due to CHS carried over from the purification.

### Achieving single-molecule resolution

The number of proteins per liposome was estimated using the Poisson distribution, as described previously^29,72^. Briefly, the number of lipid molecules (*L*) forming a vesicle is:

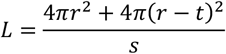

where *r* is the radius of the liposomes, *t* is the thickness of the bilayer, and *s* is the surface area of the lipid headgroups.

The number of lipids (*l*) and the number of proteins (*p*) in a given preparation can be calculated using Avogadro’s constant *N*_*A*_:

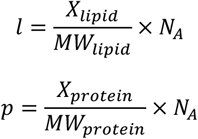

where *X* is the amount of lipid or protein, and *MW* is the molecular weight.

The number of liposomes (*N*_*liposome*_) formed in a given preparation is therefore:

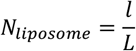

To determine the typical size distribution of the vesicles, we used cryoEM to image POPC:POPS (3:1) liposomes with 33% CHS (**ED Fig. 14a-b**). Imaging revealed two populations of vesicles: one with a diameter of 95.98 nm (87%) and the other with 38.87 nm (13%), as determined by measuring vesicle circumference in ImageJ ^73^ (**ED Fig. 14a-b**). Assuming bilayer thickness *t* is 41 Å ^74^, lipid headgroup surface area *s* is 60 Å^75^, the number of lipids needed to form a 96 nm (*L*_*96*_) or a 39 nm (*L*_*39*_) liposome would be 88607 and 12932, respectively; and the resulting number of liposomes formed (*N*_*total*_ = *N*_96_ + *N*_39_) in a given preparation (4 mg lipids per ml, 1 ml) would be 9.3 × 10^13^. Using 738 Da as an average molecular weight for lipids and 178.2 kDa for an EAAT1_fret_ trimer, assuming a ∼30% estimated reconstitution efficiency (**ED Fig. 14c-e**), about 4 × 10^13^ EAAT1_fret_ trimers were reconstituted at a PLR of 1:100 (*N*_*EAAT1*_).

Assuming the probability of liposomes containing *i* number of EAAT1fret transport follows the Poisson distribution:

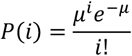

where *μ* = *N*_*EAAT*1_/*N*_*lipsome*_. The PLR of 1:100 in our reconstitution (*PLR* = *X*_*protein*_/*X*_*lipids*_) is expected to yield 65% empty liposomes. Of the 35% liposomes with EAAT1_fret_ reconstituted, 80% are expected to have one EAAT1_fret_ trimer, 17% to have two, and 3% to have three or more.

Assuming the encapsulation of the Asp/Glu sensor ccPEB1a-Y198F into liposomes also follows the Poisson model, *μ* will now be a function of the internal volume *V* of liposomes:

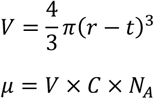

where *C* is the external concentration of ccPEB1a-Y198F. Using a *C* of 0.6 μM in our preparation, the probability of encapsulating 0, or 1 or more ccPEB1a-Y198F for the 39 nm liposomes is 99.4% and 0.6%, respectively; the probability of encapsulating 0, 1, 2 or more ccPEB1a-Y198F for the 96 nm liposomes is 87.9%, 11.3%, and 0.8%, respectively.

Taken together, many vesicles do not contain either EAAT1_fret_ or PEB1a or both. However, these vesicles are either not immobilized in the imaging chambers because they lack biotin-labeled EAAT1_fret_ or are not visible because they lack fluorescently labeled proteins. Most of the immobilized vesicles that yield FRET trajectories contain only one trimeric EAAT1_fret_ and only one ccPEB1a-Y198F for the transport assay.

### Single-molecule dynamics assay

The proteoliposomes containing labeled EAAT1_fret_ were exchanged into the appropriate buffer using PD MiniTrap G-25 columns (*Cytiva*) and subjected to three freeze-thaw cycles to replace the internal buffer. The proteoliposomes were extruded 11 times through 100-nm-pore membranes before imaging.

All imaging experiments were performed at 21 °C on a previously described home-built prism-based TIRF microscope constructed around a Nikon Eclipse Ti inverted microscope body ^26,45,76,77^. Microfluidic imaging chambers (*Finkenbeiner*) were passivated with EZ-Link™ NHS-PEG_4_-Biotin (*Thermo Scientific*) and mPEG-SPA-1000 (*Laysan Bio*) as previously described ^26,27,29,45^. After passivation, the microfluidic channels were incubated with 0.8 μM neutravidin (*Invitrogen*), 2 μM 25-nucleotide DNA duplex (IDT), and 2 μM BSA (EM-2930, VWR) in T50 buffer (10 mM Tris, pH 7.5, and 50 mM KCl) for 10 min, then thoroughly rinsed with T50 buffer and experimental buffer. EAAT1_fret_ proteoliposomes were immobilized by slowly flowing them over the channel; excess sample was removed by washing with experimental buffers containing 50 mM HEPES/Tris, pH 7.4, 2.5% glycerol, and, as appropriate: 200 mM NaCl with various concentrations of L-aspartate or L-glutamate (NaCl buffer); 200 mM KCl (KCl buffer); or 200 mM Tris/HCl (TrisCl buffer). All buffers were supplemented with an oxygen-scavenging system composed of 2 mM protocatechuic acid and 50 nM protocatechuate 3,4-dioxygenase, as described previously ^77^.

Immobilized proteoliposomes were illuminated with a 532-nm laser (*Laser Quantum*). LD555 and LD655 fluorescence signals were separated using a T635lpxr dichroic filter (*Chroma*) mounted in a MultiCam apparatus (*Cairn*). Imaging data were acquired using scientific complementary metal–oxide–semiconductor (sCMOS) cameras ORCA®-Fusion BT (*Hamamatsu*). The smFRET movies were recorded under equilibrium conditions (same ionic conditions on both sides of the proteoliposomes) and under real-time-initiated flux conditions. For uptake conditions, proteoliposomes were prepared in a buffer containing 200 mM KCl, and perfused with a buffer containing 200 mM NaCl and varying concentrations of L-aspartate or L-glutamate approximately 2 s after the start of the recordings. For efflux conditions, proteoliposomes were prepared in buffers containing 200 mM NaCl and an appropriate concentration of L-aspartate or L-glutamate, then perfused with 200 mM KCl as above. All recordings were performed with a 100-ms integration time at 100 mW laser power, except where noted, when a 200-ms integration time with 50 mW laser power was used.

### Analysis of EAAT_fret_ dynamics

Traces were analyzed using Spartan software ^78^ implemented in MATLAB (*MathWorks*). FRET trajectories (*E*_*FRET*_) were calculated as described previously ^76,79^ from LD555 and LD655 intensities, *I*_*D*_ and *I*_*A*_, respectively, as *E*_*FRET*_ = *I*_*A*_/(*I*_*D*_ + *I*_*A*_). Trajectories were corrected as described previously ^76,79^ for spectral cross-talk, unequal apparent brightness of donor and acceptor fluorophores, and acceptor direct excitation, and automatically preprocessed to exclude those lasting ≤30 frames and with a signal-to-noise ratio ≤8. Traces with multiple photobleaching events, indicative of multiple transporters in the liposomes or incorrect labeling, inconsistent total fluorescence intensity, or lack of anticorrelated fluorescence intensity were discarded. Corrected FRET values (E) were then used for estimating inter-dye distances (R) using 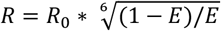 and an R_0_ of 62 Å^80^.

For each condition, smFRET traces, corresponding to single EAAT1_fret_ transporters, were recorded in 30-50 movies across three independent setups, with proteins purified from at least two different transfections. All traces (*total* molecules) were sorted into *2-active, 1-active*, 2-*pausing*, and *static* before idealizing. *E*_*FRET*_ trajectories belonging to *2-active, 1-active*, and 2-*pausing* were idealized separately using a five-state model in Spartan^76^, with the following mean (± S.D.) starting values: 0 (0.04), 0.23 (0.04), 0.43 (0.06), 0.59 (0.06), 0.78 (0.04). Traces showing modal switching were sorted according to their predominant behavior. Dwelltime distributions and transition density plots were generated as previously described ^78^ and plotted in MATLAB (*MathWorks*). Dwell-time histograms were normalized by dividing the number of dwells in each duration bin by the total imaging time, yielding the number of dwells per second. This approach ^79^ ensures that the histograms do not overemphasize rare transitions, that the y-axis height relates to transition frequency, and that it is minimally affected by variations in molecule number and photobleaching. The resulting dwell-time histograms were integrated to obtain the populations of each FRET state, which were used to estimate the population of the three conformational states, OFS, intS, and IFS using the following relationships between populations of conformational states and FRET states (FRET0.8, FRET0.6, FRET0.4, and FRET0.2), where IF/IF state contributes to FRET0.8, IF/int state to FRET0.6, OF/IF and int/int states to FRET0.4, and OF/OF and OF/int to FRET0.2:

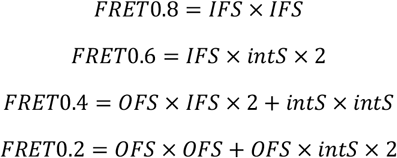

The resulting populations of OFS, intS, and IFS were used to back-calculate the expected populations of the 0.8, 0.6, 0.4, and 0.2 FRET states, assuming that protomers behave independently of each other.

The mean transition rates of transporters were averaged from the transition rates of individual transporters. The dwell-time survival plots were fitted to bi- or tri-exponential decay functions, and the histograms were fitted to transformed probability density functions ^81^.

### Single-transporter activity assays

Single-transporter assays were performed on the TIRF microscope using the same experimental setup as the single-molecule dynamics assays described above. Proteoliposomes containing reconstituted biotin-PEG_11_-labeled EAAT1_fret_ and encapsulated LD555/LD655-labeled ccPEB1a-Y198F were surface-immobilized in the microfluidic chamber. smFRET movies were recorded with 200 ms integration time and 50 mW laser power to capture real-time transport events, or with 100 ms integration time and 100 mW laser power to monitor the equilibrated state. Vesicles were prepared in 200 mM NaCl and 100 µM L-aspartate. To confirm that the proteoliposomes were not leaky to L-aspartate, control recordings were first taken in a buffer containing 200 mM NaCl, followed by efflux initiated by perfusing a buffer containing 200 mM KCl. To measure uptake, the same vesicles emptied during efflux were perfused first with a buffer containing 200 mM KCl and 10 µM L-aspartate (to test for leaks) and then with a buffer containing 200 mM NaCl and 10 µM L-aspartate. Vesicles showing altered FRET efficiency in the control buffers, indicative of L-aspartate leak, were excluded from further analysis. In all experiments, the appropriate buffers were perfused 2-3 s after the start of the recording. The number of aspartate molecules *N*_*Asp*_ in a 100 nm diameter vesicle is calculated as a function of the internal volume *V*:

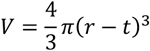

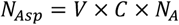

where *C* is the internal concentration of aspartate, *r* is the radius of the liposomes, *t* is the thickness of the bilayer, and *N*_*A*_ is Avogadro’s constant.

### Radioactive transport assay in HEK293 cells

Cell-based transport uptake assays were performed as described previously ^31^. Cells were collected 50 hr post-transfection, resuspended at 50 × 10^6^ cells ml^−1^ in resting buffer (11 mM HEPES/Tris-base, pH 7.4, 140 mM ChCl, 4.7 mM KCl, 2.5 mM CaCl_2_, 1.2 mM MgCl_2_, and 10 mM D-glucose), and used immediately. Substrate transport was assayed at 37 °C. The resuspended cells were diluted 5-fold into reaction buffer (11 mM HEPES/Tris-base, pH 7.4, 140 mM NaCl, 4.7 mM KCl, 2.5 mM CaCl_2_, 1.2 mM MgCl_2_, 10 mM D-glucose) or resting buffer for background estimation. Background was also measured in untransfected cells that underwent the same procedures. Uptake reactions were initiated by adding freshly prepared L-aspartate solution to a final concentration of 19.5 µM cold L-aspartate and 0.5 µM L-[^3^H]-Asp (aspartic acid, L-[2,3-^3^H], *PerkinElmer*). At each time point, aliquots of the uptake reaction were diluted 20-fold into ice-cold resting buffer to halt transport. Samples were immediately filtered through 0.8 μm nitrocellulose filters (*Millipore*) and washed three times with 3 ml of ice-cold resting buffer. The washed filter membranes were transferred into scintillation vials, and the retained radioactivity was measured using the LS-6500 scintillation counter (*Beckman Coulter*). Each time-course experiment was fitted to the one-phase association equation (*GraphPad Prism*):

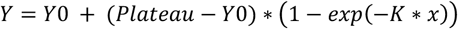

and each time point was normalized to the maximum from the fit. Five to nine independent time-course experiments with cells from two to three different transfections were performed per mutant or EAAT1_fret_.

## Data availability

Atomic coordinates and EM densities have been deposited in PDB and EMDB for the following structures: NaAsp bound IFS tight (9ZDT, EMD-74073), NaAsp bound IFS loose (9ZDZ, EMD-74079), NaAsp bound IFS loose* (9ZQK, EMD-74568), V309G NaAsp bound IFS loose (9ZQL, EMD-74569), V309G NaAsp bound IFS loose* (9ZQM, EMD-74570), Y288L NaAsp bound IFS tight (9ZKB, EMD-74360), OFS apo (9ZQH, EMD-74565), IFS apo (9ZQJ, EMD-74567), NaAsp bound OFS free (9ZQW, EMD-74583), NaAsp bound OFS wrapped (9ZR8, EMD-74596), NaAsp bound intS (9ZU9, EMD-7799), NaAsp bound IFS shift (9ZU2, EMD-74788), TFB-TBOA bound IFS loose (9ZU9, EMD-74799), TFB-TBOA bound IFS loose* (9ZUE, EMD-74808), P290R NaAsp bound IFS straight (10BN, EMD-75046), P290R NaAsp bound IFS tight (10BQ, EMD-75051), P290R NaAsp bound IFS tight* (10BP, EMD-75052), P290R NaAsp bound IFS straight* (10BP, EMD-75050), P259R E3 IFS open (10CI, EMD-75063), P259R E3 NaAsp bound IFS closed (10CJ, EMD-75064), P259R E3 NaAsp bound cholesterol bound IFS closed* (10CO, EMD-75069). Extended Figure 9 has all raw data for each condition of the smFRET recordings. All other plasmids and data are available upon request.

## Acknowledgements

We thank Dr. Scott C. Blanchard for assistance with smFRET instruments. We thank Drs. Xiaoyu Wang, Krishna Reddy, Biao Qiu, Andréia C.K. Mortensen, Ole Mortensen, and Katelyn Reeb for helpful discussions. We thank Drs. Zhang Feng and Philipp Schmidpeter for helpful discussions on nanodisc studies. We thank Vishnu Ghani and Dr. Chieh-Chin Li for helpful discussions on developing in-house scripts for MATLAB-based analysis. We thank the Scientific Computing Unit at Weill Cornell Medical College for the maintenance and support of computational resources. We thank graduate students Lujia Gao and Alexander Earsley for their assistance during their rotation. We thank Drs. Carl Fluck at Weill Cornell Cryo-EM Core Facility, Huihui Kuang, Bing Wang, and William Rice at New York University Langone cryo-EM laboratory, Robert Grassucci at the Columbia University Medical Center Cryo-EM core, Rui Yan and Xiaowei Zhao from Janelia cryoEM facility for assistance with cryoEM data collection. We thank Lesley Anson, Vishnu Ghani, and Alexander Earsley for helpful comments and proofreading the manuscript.

The work was supported by Charles H. Revson Senior Fellowship in Biomedical Science CHRF 24-34 (to Q. Wu), American Heart Association Postdoctoral Fellowship 909426 (to Q. Wu), and National Institutes of Health NIH grants R01NS064357, R37NS085318, and R37NS134865 (to O. Boudker). O. Boudker is an investigator of the Howard Hughes Medical Institute.

## Author contributions

Q.W., D.C., and O.B. conceived the project. Q.W. conceptualized, designed, and performed all smFRET, cryoEM, and activity experiments and analyzed the data. Q.W. and O.B. interpreted the data and directed the project. D.C. designed and performed foundational preliminary experiments. J.C.C. and N.R. developed protein expression constructs and optimized protein expression and purification. G.G.G. assisted with smFRET microscope optimization. Y.H. assisted with NMR experiments. Q.W., J.C.C., N.R., and O.B. participated in regular discussions. Q.W. and O.B. wrote the manuscript. All authors reviewed and edited the manuscript.

