## Supplementary material for "Multimodal dynamics control activity of a glial glutamate transporter": Supplymentary Figures

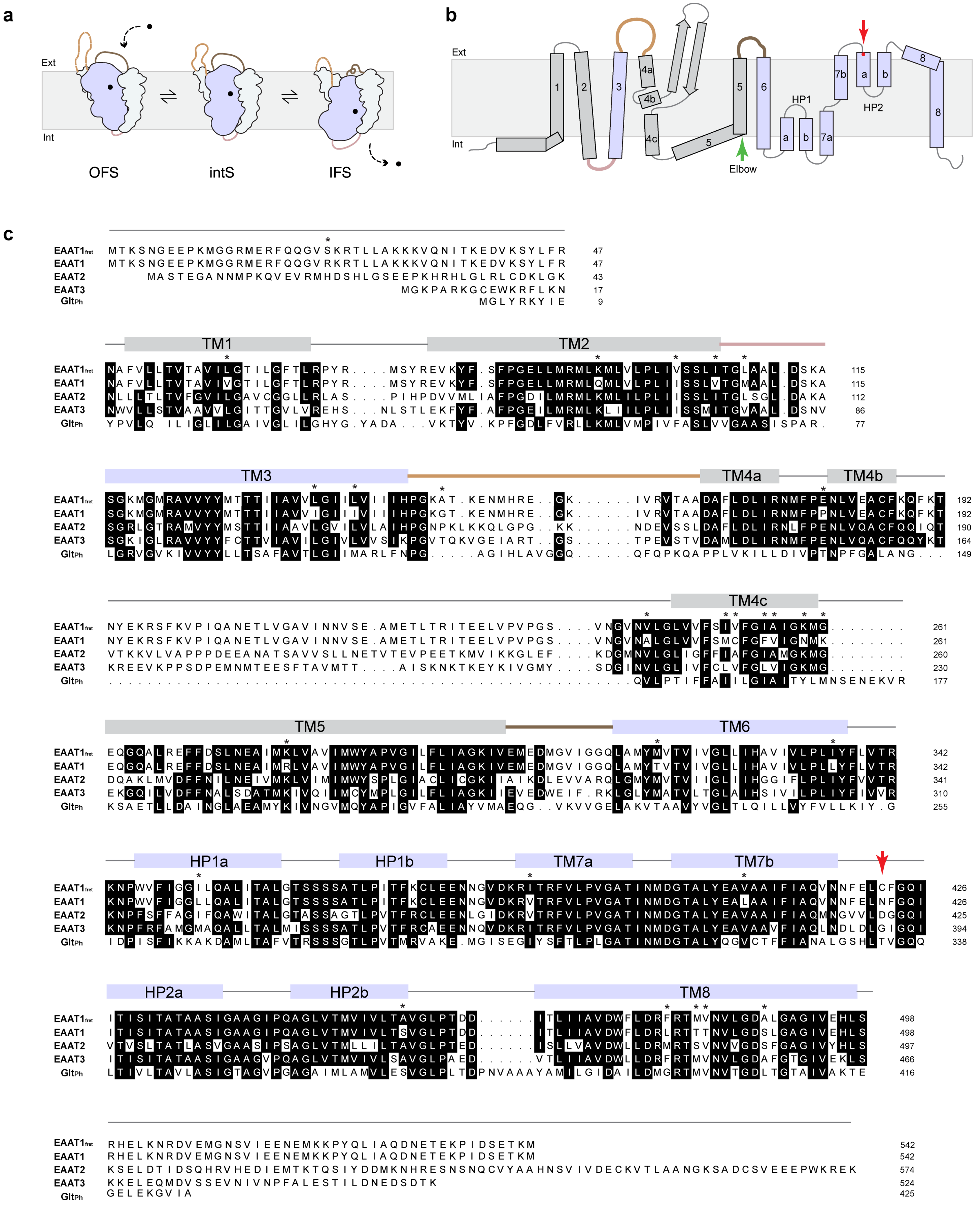


Extended Data Figure 1. Thermally stabilized human EAAT1 ($EAAT1_{fret}$).

**a**, Alternating-access elevator mechanism of glutamate transporters, with a protomer transitioning through outward-facing (OFS), intermediate (intS), and inward-facing (IFS) states. The protomer comprises the stationary scaffold (gray) and mobile transport (lavender) domains, connected by hinges and loops (brown). Substrate and coupled ions (black dot) bind within the transporter domain and are translocated from the extracellular to the intracellular side of the membrane (light gray area). **b**, TM topology of EAAT1$_{fret}$ showing three hinges/loops linking the transport domain to the scaffold (color scheme as in **a**). The TM5 elbow and the location of the N422C mutation for fluorescent labeling are marked by green and red arrows, respectively. **c**, The sequence alignment of the EAAT1$_{fret}$ construct, derived from thermally stabilized cocoEAAT1, with human EAAT2 and 3, and archaeal homolog Glt$_{Ph}$. The N422C mutation is marked by a red arrow. Conserved residues are shaded black; asterisks denote mutations relative to human EAAT1. Colored bars indicate transmembrane segments (TMs) and reentrant helical hairpins (HPs).

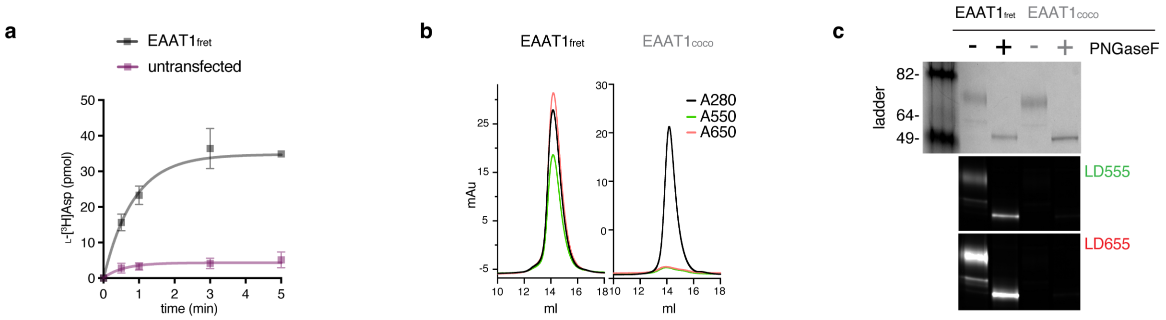


**Extended Data Figure 2. EAAT1$_{fret}$ is a functional, monodisperse protein specifically labeled with LD555 and LD655 donor and acceptor fluorescent dyes.**

**a,** EAAT1$_{fret}$ heterologously expressed in HEK293 cells supports robust L-[$^3$H]Asp uptake. **b,** Purified protein labeled with a mixture of biotin-(PEG)$_{11}$-maleimide and maleimide derivatives of LD555 and LD655 elutes as a single peak during size-exclusion chromatography (SEC), monitored by absorbance at 280 (black line), 550 (green), and 650 (red) nm. Labeling efficiency was 57.0 ± 0.8% and 57.1 ± 1.5% for LD555 and LD655, respectively. In contrast, parent cocoEAAT1 lacking the N422C mutation showed no significant labeling. **c,** SDS-PAGE analysis of labeled EAAT1$_{fret}$ and cocoEAAT1. PNGase F treatment confirms that the protein is glycosylated. Coomassie staining is shown on top, with fluorescent imaging using LD555- and LD655-specific filters below.

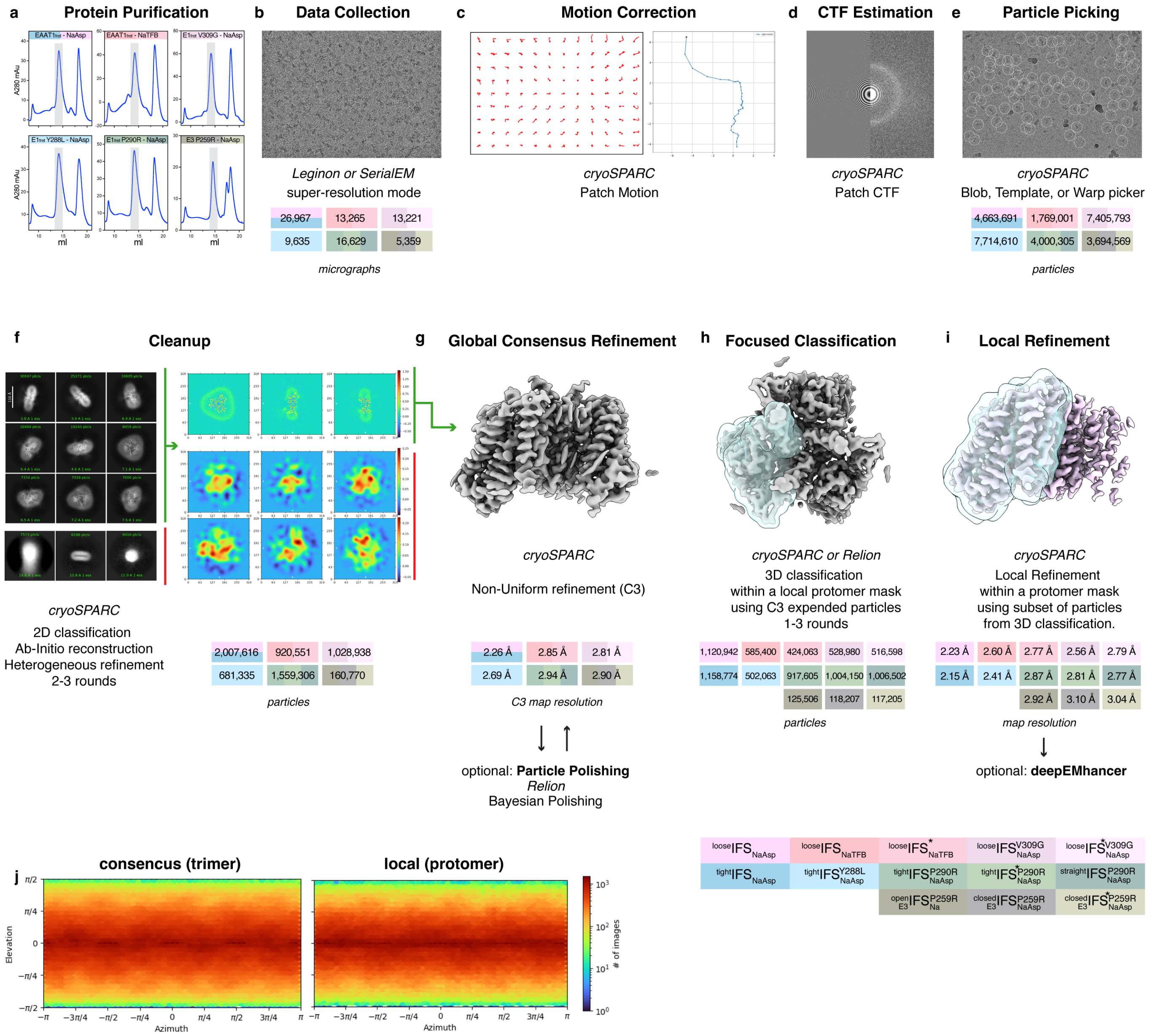


### Extended Data Figure 3. CryoEM data processing in detergent.

Data are shown for EAAT1$_{fret}$ and mutants. Colors denote protein preparations and structural classes resolved after processing. **a**, SEC profiles of EAAT1$_{fret}$ in 200 mM Na$^+$ and 1 mM L-aspartate (EAAT1$_{fret}$-NaAsp); EAAT1$_{fret}$ purified in buffer containing 200 mM Na$^+$ and 1 mM L-aspartate, then exchanged into and frozen with 200 mM Na$^+$ and 0.4 mM TFB-TBOA (EAAT1$_{fret}$-NaTFB); V309G (E1$_{fret}$ V309-NaAsp); Y288L (E1$_{fret}$ Y288L-NaAsp); P290R (E1$_{fret}$ P290R-NaAsp); and EAAT3 P259R (E3 P259R-NaAsp). All proteins were purified in 0.48% DDS and 0.12% CHS. Grey shading marks fractions used for cryo-EM; the later-eluting peak corresponds to cleaved GFP. **b**, Representative cryoEM micrograph; movie numbers per dataset are shown in colored boxes. **c**, Motion correction (cryoSPARC). **d**, CTF estimation. **e**, Particle picking; particle numbers shown in colored boxes. **f**, Particle cleaning by 2D classification and heterogeneous refinement; green and red bars indicate selected and rejected classes, respectively; retained particle numbers shown in colored boxes. **g**, Nonuniform refinement with C3 symmetry; map resolutions shown in colored boxes. **h-i**, Symmetry expansion and 3D classification (**h**), followed by local refinement with a protomer mask (**i**); particle numbers (**h**) and map resolutions (**i**) shown in colored boxes. **j**, Representative heat maps of particle angular distributions.

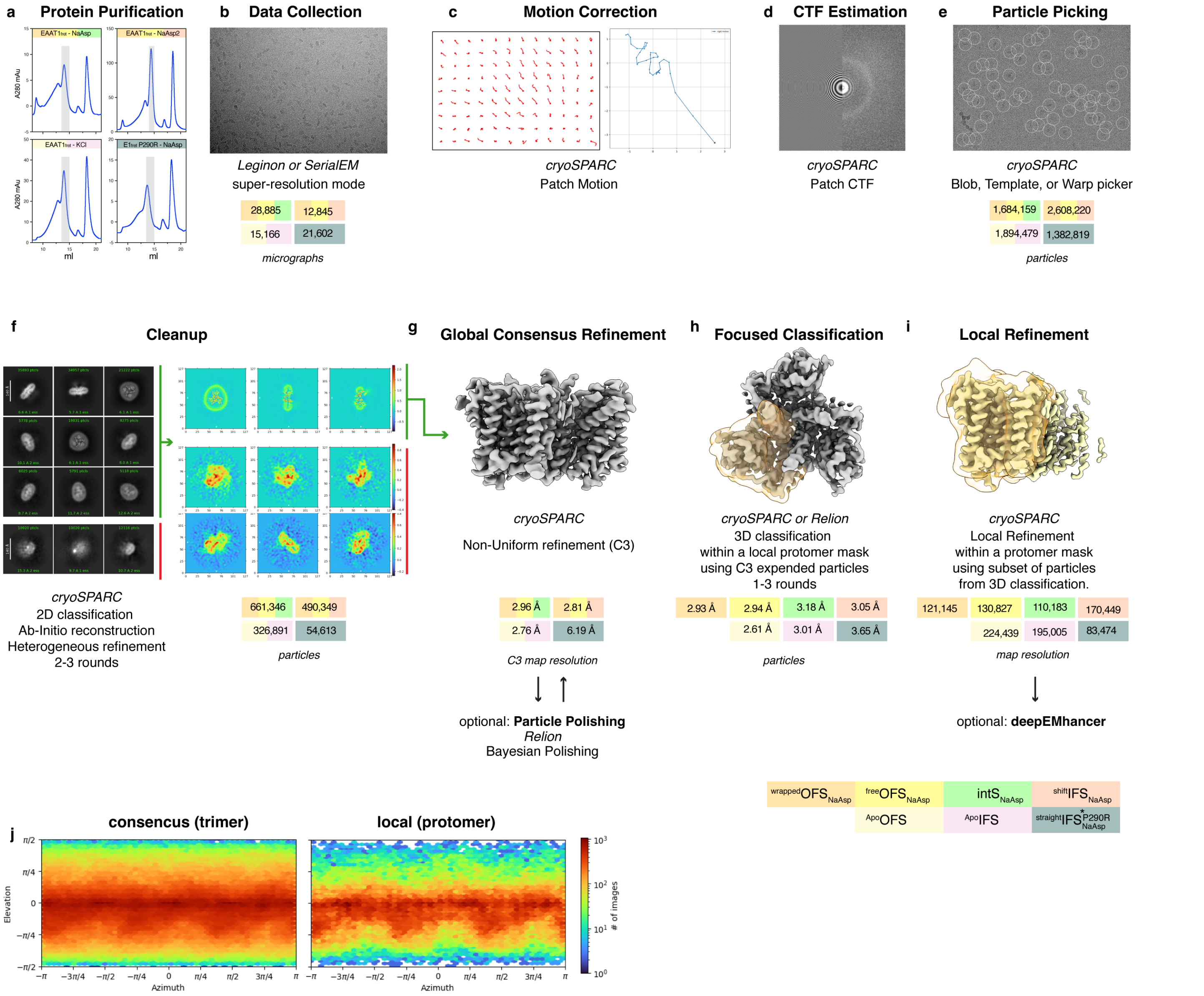


#### Extended Data Figure 4. CryoEM data processing in nanodiscs.

Data are shown for EAAT1$_{fret}$ and mutant proteins. The color scheme reflects the protein preparation and the structural classes resolved at the end of the processing pipeline. **a**, SEC profiles of two preparations of MSP2N2-reconstituted EAAT1$_{fret}$ in 200 mM $Na^+$ and 1 mM Asp (EAAT1$_{fret}$-NaAsp and EAAT1$_{fret}$-NaAsp2); EAAT1$_{fret}$ in 200 mM $K^+$ (EAAT1$_{fret}$-KCl); and EAAT1$_{fret}$ P290R in 200 mM $Na^+$ and 1 mM Asp (E1$_{fret}$ P290R-NaAsp). Because the first preparation of EAAT1$_{fret}$ in NaAsp was low-yielding, a second preparation was performed with more cells. **b**, Representative cryoEM micrograph; movie numbers per dataset are shown in colored boxes. **c**, Motion correction (cryoSPARC). **d**, CTF estimation. **e**, Particle picking; particle numbers shown in colored boxes. **f**, Particle cleaning by 2D classification and heterogeneous refinement; green and red bars indicate selected and rejected classes, respectively; retained particle numbers shown in colored boxes. **g**, Nonuniform refinement with C3 symmetry; map resolutions shown in colored boxes. **h-i**, Symmetry expansion and 3D classification (**h**), followed by local refinement with a protomer mask (**i**); particle numbers (**h**) and map resolutions (**i**) shown in colored boxes. **j**, Representative heat maps of particle angular distributions.

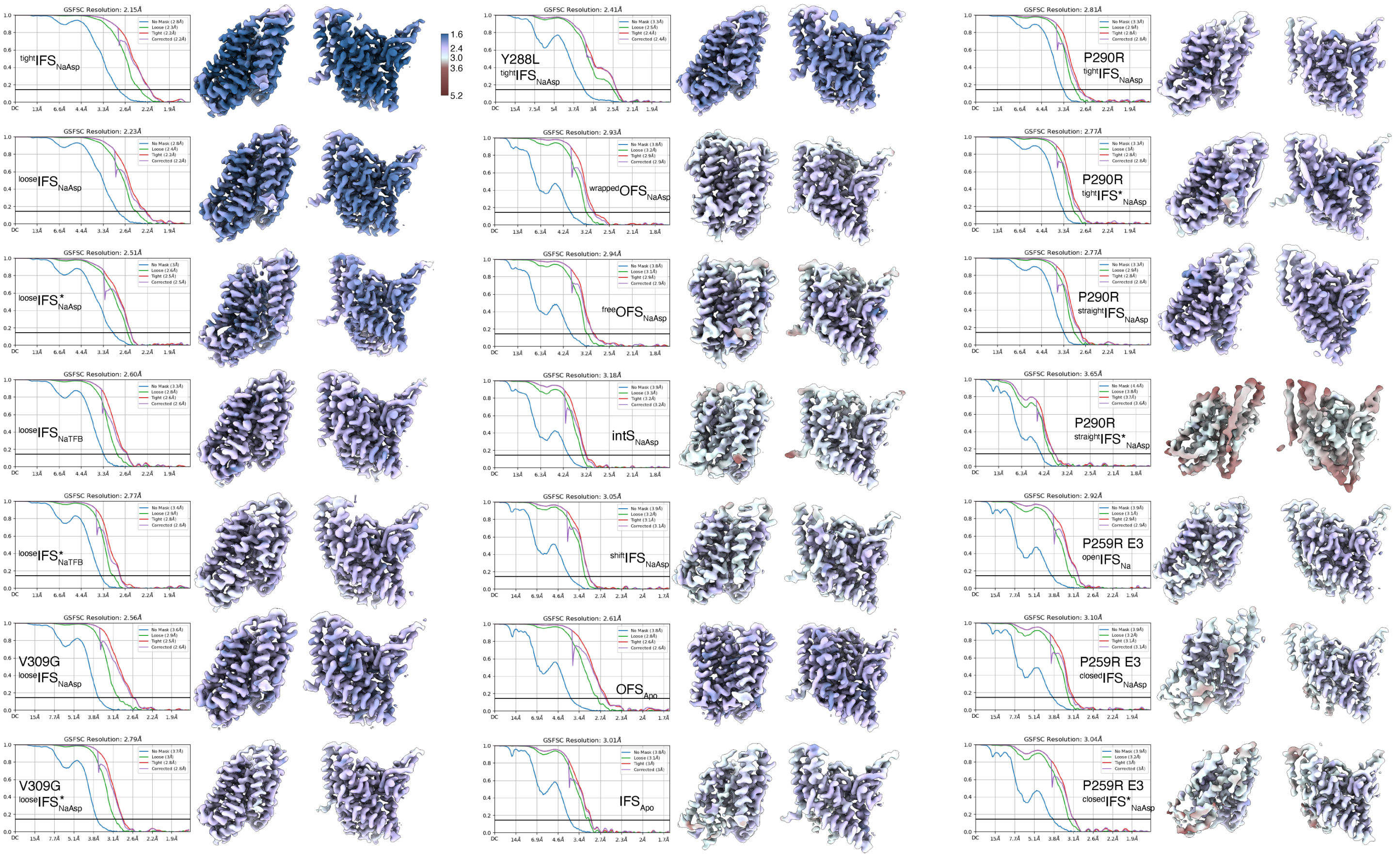


**Extended Data Fig. 5 FSC and local resolution maps for all structures.**

Structures were determined from the datasets in Extended Data Figs. 3 and 4. EAAT1$_{fret}$ - NaAsp (detergent): $^{tight}IFS_{NaAsp}$, $^{loose}IFS_{NaAsp}$, $^{loose}IFS'_{NaAsp}$; EAAT1$_{fret}$-NaTFB (detergent): $^{loose}IFS_{NaTFB}$, $^{loose}IFS'_{NaTFB}$; E1$_{fret}$ V309G-NaAsp (detergent): V309G $^{loose}IFS_{NaAsp}$, V309G $^{loose}IFS'_{NaAsp}$; E1$_{fret}$ Y288L-NaAsp (detergent): Y288L $^{tight}IFS_{NaAsp}$; EAAT1$_{fret}$-NaAsp (nanodiscs): $^{wrapped}OFS_{NaAsp}$, $^{free}OFS_{NaAsp}$, $IntS_{NaAsp}$; EAAT1$_{fret}$-NaAsp (nanodiscs) 2: $^{shift}IFS_{NaAsp}$ ($^{wrapped}OFS_{NaAsp}$ and $^{free}OFS_{NaAsp}$ were also observed in this dataset at similar resolution); EAAT1$_{fret}$-KCl (nanodiscs): $OFS_{Apo}$, $IFS_{Apo}$; E1$_{fret}$ P290R-NaAsp (detergent): P290R $^{tight}IFS'_{NaAsp}$, P290R $^{tight}IFS'_{NaAsp}$, P290R $^{straight}IFS_{NaAsp}$; E1$_{fret}$ P290R-NaAsp (nanodiscs): P290R $^{straight}IFS'_{NaAsp}$; E3 P259R-NaAsp (detergent): P259R E3 $^{open}IFS_{Na}$, P259R E3 $^{closed}IFS_{NaAsp}$, P259R E3 $^{CLR}IFS_{NaAsp}$. TFB-TBOA was bound in two distinct poses ($^{loose}IFS_{NaTFB}$, $^{loose}IFS'_{NaTFB}$), and we think that both structures might resemble $Na^{+}$-bound states as seen in other family members [2]. The local resolution scale bar ranges from 1.6 Å (dark blue) to 5.2 Å (deep red).

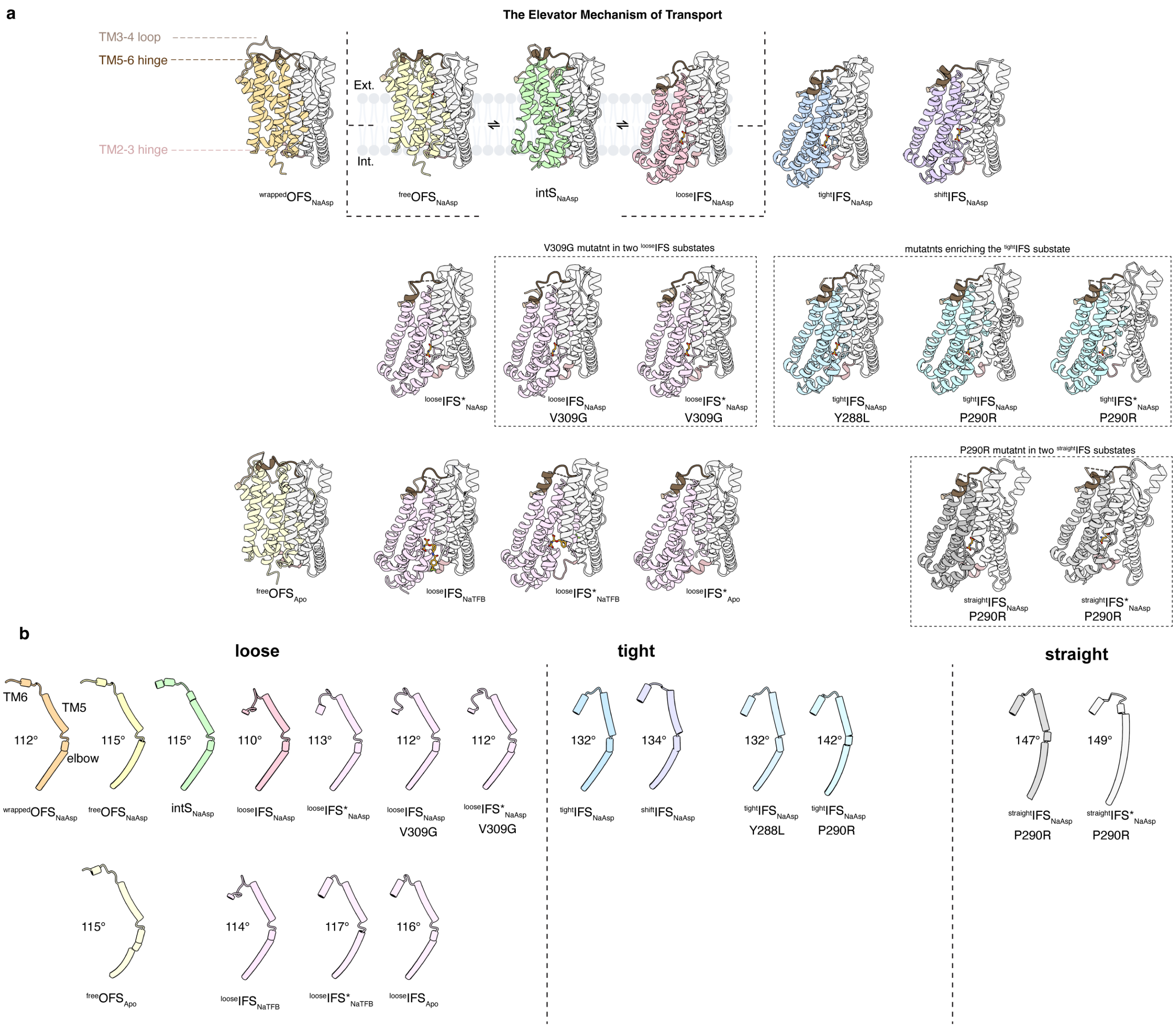


#### Extended Data Fig. 6 Snapshots of the transport cycle with highlighted elbow structures.

**a**, CryoEM structures are shown in cartoon representation and arranged along the envisioned transport cycle. The three central structures, $^{free}$OFS$_{NaAsp}$, intS$_{NaAsp}$, and $^{loose}$IFS$_{NaAsp}$, depict movement of the transport domain (colored) relative to the scaffold domain (white) across the membrane. $^{wrapped}$OFS$_{NaAsp}$ and $^{tight}$IFS$_{NaAsp}$ are proposed to be off-cycle states; entry into these states might lead to pausing and slow turnover. $^{shift}$IFS$_{NaAsp}$ is a structural class with an elbow structure similar to $^{tight}$IFS$_{NaAsp}$ but with distortions in the scaffold domain and an associated shift in the transport domain; non-protein density, possibly a lipid molecule or CHS, is inserted between scaffold TM4 and TM5. Other structural classes observed for EAAT1$_{fret}$ and mutants, with the transporter bound to $Na^+$/Asp, $Na^+$/TFB-TBOA, or in the *apo* state, are shown below, with transport domains colored yellow for OFS, pink for $^{loose}$IFS, blue for $^{tight}$IFS, and gray for $^{straight}$IFS of the P290R mutant. Hinges and loops between the scaffold and transport domains are colored in shades of brown. **b**, Tube representation of TM5 and the beginning of TM6, highlighting differences in the elbow bend. A different structure of the TM5-6 hinge is observed in the substate $^{loose}$IFS*, with TM5 bent to a similar degree as in $^{loose}$IFS. Estimated angles are shown next to the cartoons. Color scheme is as in panel **a**.

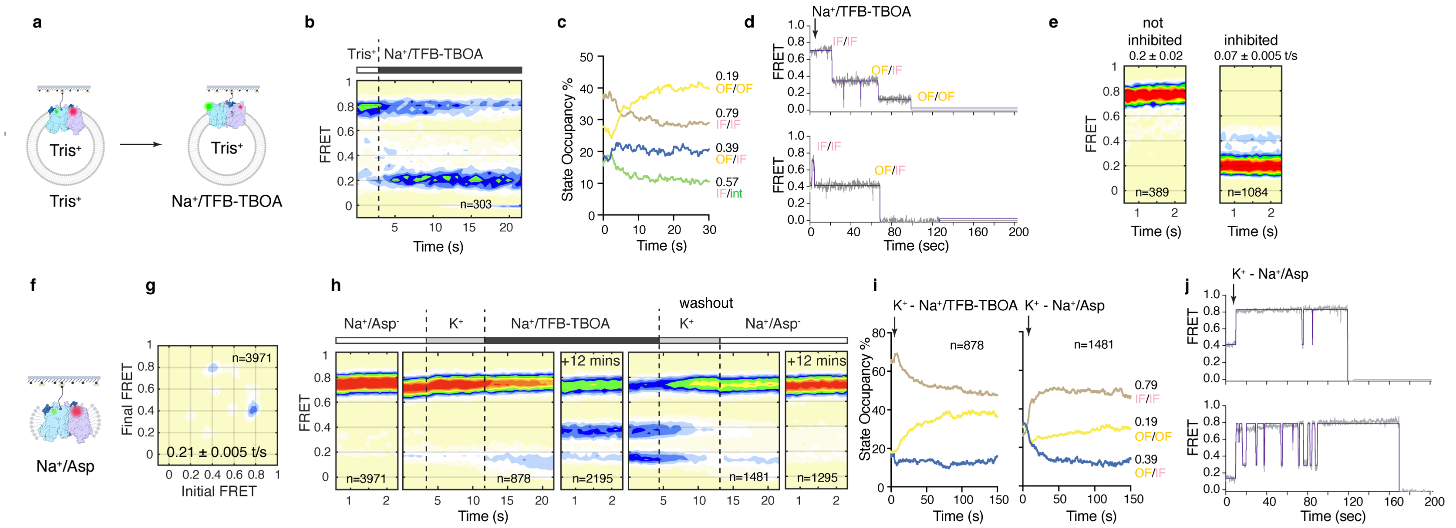

**Extended Data Fig. 7. Ions, substrate, and TFB-TBOA blocker affect EAAT1$_{fret}$ dynamics.**

**a**, Proteoliposomes equilibrated in *apo* Tris buffer (no alkali ions) and perfused in real time with 200 mM NaCl and 400 µM TFB-TBOA ~2 s after recording onset. **b**, FRET state population plots from the first 22 s of recordings; molecule numbers (*n*) are indicated. Bars denote the external solution: *apo* Tris (white) or $Na^+$/TFB-TBOA (black). **c**, Time course of averaged state occupancies after $Na^+$/TFB-TBOA application (data from b). FRET states and corresponding conformations are indicated. **d**, Example raw smFRET traces; an arrow marks the injection time. Raw data, gray; idealized traces, purple. **e**, Population plots for molecules that did not change (not inhibited) or did change (inhibited) FRET state 12 min following $Na^+$/TFB-TBOA perfusion; transition frequencies shown above. **f-i**, TFB-TBOA binding and unbinding in DDS. Protein equilibrated in 200 mM NaCl and 1 mM Asp (**f**) shows little dynamics (**g**). After recording in $Na^+$/Asp (**h**), a new recording was initiated; ~2 s later, the sample was perfused with 200 mM KCl for ~8 s to remove substrate, then with 200 mM NaCl and 400 µM TFB-TBOA. After 12 min of equilibration, an additional movie was recorded. For washout, a new recording was initiated, and proteoliposomes were perfused with KCl, then $Na^+$/Asp. Bars denote external solution: $Na^+$/Asp (white), $K^+$ (gray), or $Na^+$/TFB-TBOA (black). **i**, Averaged occupancy time courses after KCl followed by $Na^+$/TFB-TBOA (left) or $Na^+$/Asp (right); injections marked by arrows. **j**, Example raw smFRET traces of TFB-TBOA washout; raw data, gray; idealized traces, purple.

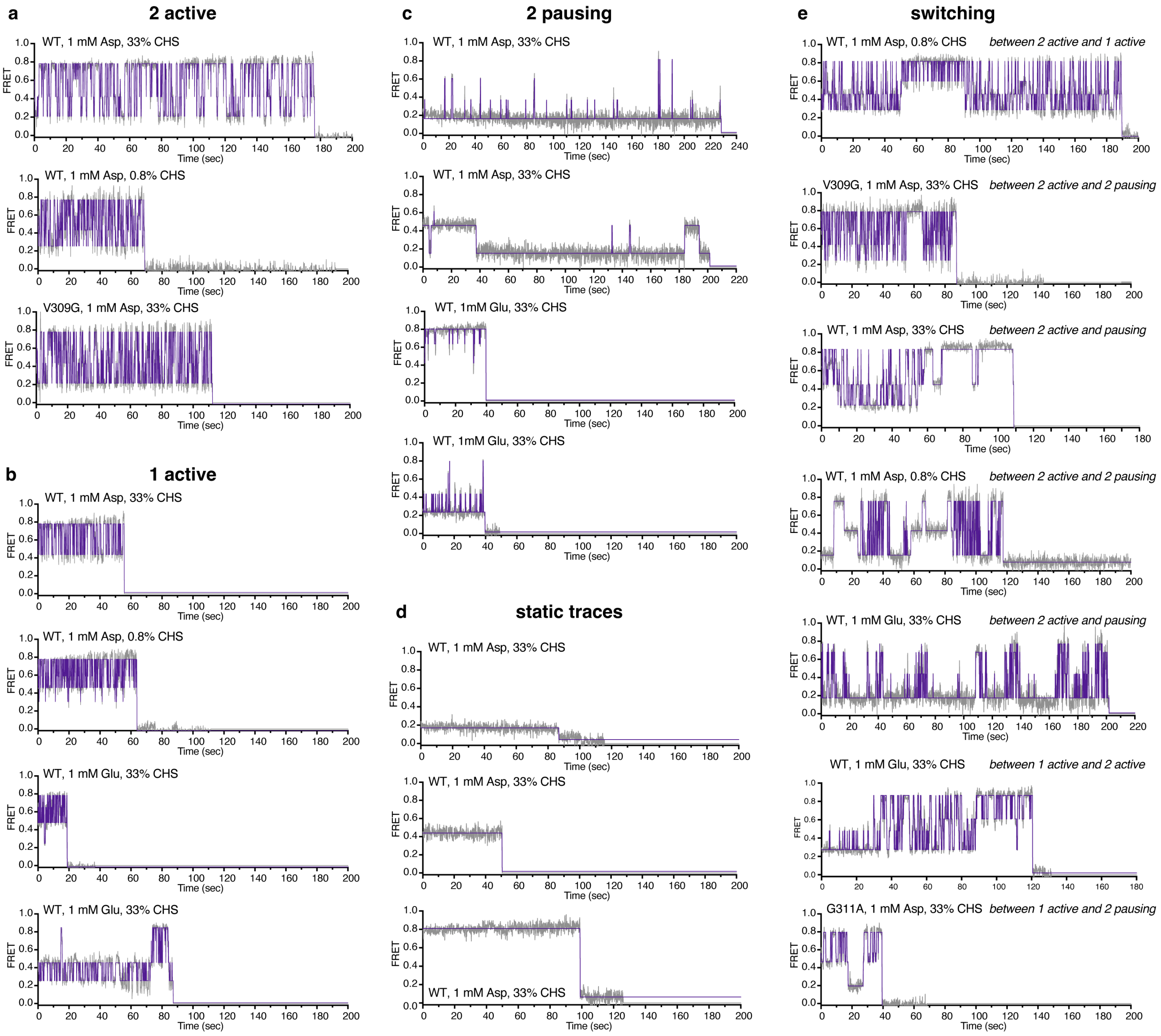


**Extended Data Fig. 8. Example smFRET traces.**

**a-e,** Representative traces of EAAT1$_{fret}$ in liposomes prepared in 200 mM NaCl and 1 mM substrates recorded in the same external buffer. Shown are traces for transporters with both observed protomers actively sampling OFS, intS, and IFS (**a**), one active and one pausing protomer (**b**), two pausing protomers with brief, infrequent transitions, most occurring at a frequency of less than 0.5 $s^{-1}$ (**c**), static traces (**d**), and traces switching between 2-active, 1-active, and pausing modes (**e**). Raw data are in gray, and idealized traces are in purple. Constructs, substrates, and CHS content of the liposomes are shown above the panels.

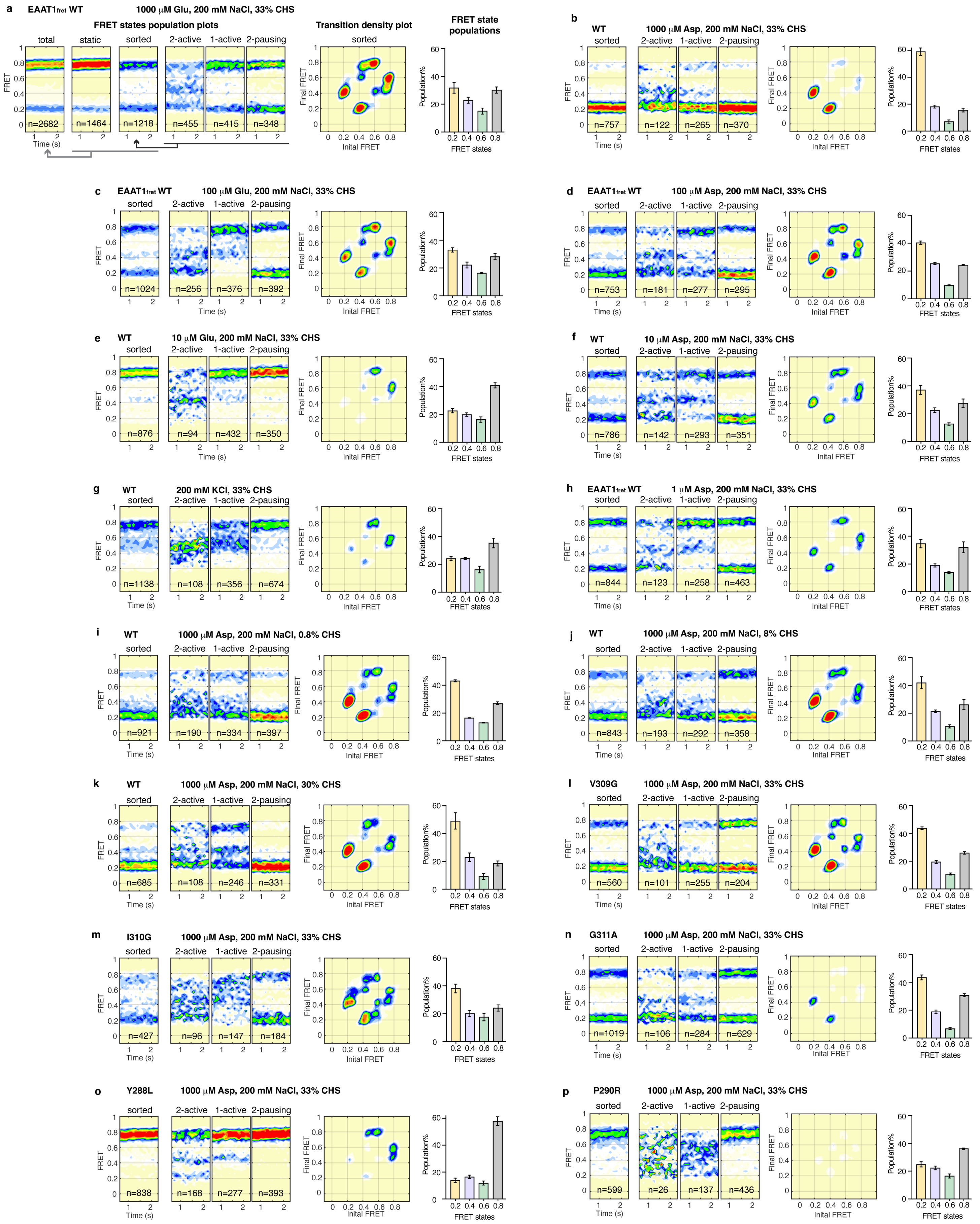
a
EAATfret WT
1000 μM Glu, 200 mM NaCl, 33% CHS
FRET states population plots
total
static
sorted
2-active
1-active
2-pausing
n=2682
n=1464
n=1218
n=455
n=415
n=348
Time (s)
FRET
Transition density plot
sorted
Final FRET
Inital FRET
FRET state populations
Population%
FRET states
b
WT
1000 μM Asp, 200 mM NaCl, 33% CHS
n=757
n=122
n=265
n=370
c
EAAT1fret WT
100 μM Glu, 200 mM NaCl, 33% CHS
n=1024
n=256
n=376
n=392
d
EAAT1fret WT
100 μM Asp, 200 mM NaCl, 33% CHS
n=753
n=181
n=277
n=295
e
WT
10 μM Glu, 200 mM NaCl, 33% CHS
n=876
n=94
n=432
n=350
f
WT
10 μM Asp, 200 mM NaCl, 33% CHS
n=786
n=142
n=293
n=351
g
WT
200 mM KCl, 33% CHS
n=1138
n=108
n=356
n=674
h
EAAT1fret WT
1 μM Asp, 200 mM NaCl, 33% CHS
n=844
n=123
n=258
n=463
i
WT
1000 μM Asp, 200 mM NaCl, 0.8% CHS
n=921
n=190
n=334
n=397
j
WT
1000 μM Asp, 200 mM NaCl, 8% CHS
n=843
n=193
n=292
n=358
k
WT
1000 μM Asp, 200 mM NaCl, 30% CHS
n=685
n=108
n=246
n=331
l
V309G
1000 μM Asp, 200 mM NaCl, 33% CHS
n=560
n=101
n=255
n=204
m
I310G
1000 μM Asp, 200 mM NaCl, 33% CHS
n=427
n=96
n=147
n=184
n
G311A
1000 μM Asp, 200 mM NaCl, 33% CHS
n=1019
n=106
n=284
n=629
o
Y288L
1000 μM Asp, 200 mM NaCl, 33% CHS
n=838
n=168
n=277
n=393
p
P290R
1000 μM Asp, 200 mM NaCl, 33% CHS
n=599
n=26
n=137
n=436

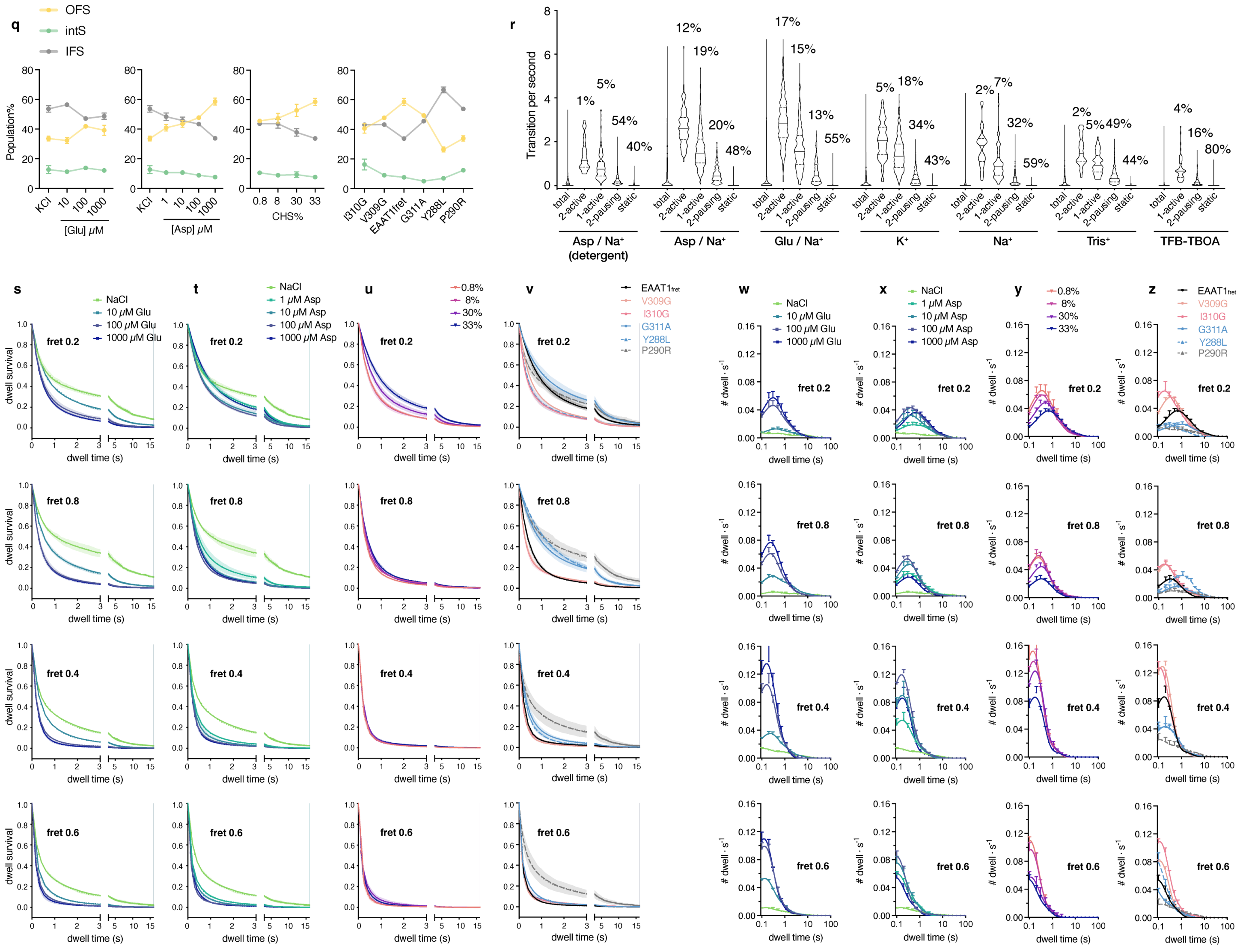


#### Extended data figure 9. Analysis of EAAT1$_{fret}$ smFRET dynamics.

**a**, smFRET analysis of EAAT1$_{fret}$ in proteoliposomes prepared in 200 mM NaCl and 1000 μM Glu. From left to right: population plots for all molecules (total), dynamic (sorted), static, 2-active, 1-active, and 2-pausing molecules; transition density plot of sorted molecules; and FRET-state populations derived from experimentally measured dwell times of sorted molecules. **b-p**, Similar analysis (total molecules are not shown) for EAAT1$_{fret}$ under varied substrate and ion conditions (**b-h**) and CHS content (**b, i-k**), and for mutants V309G (**l**), I310G (**m**), G311A (**n**), Y288L (**o**), and P290R (**p**) in 200 mM NaCl, 1000 μM Asp, and 33% CHS. Constructs and conditions are indicated above the panels. Each dataset comprises 30–50 movies from 2–3 TIRF sessions from 2–3 independent proteoliposome preparations. **q**, Comparisons of the deconvoluted populations of OFS, intS, and IFS across various Glu or Asp concentrations, CHS content, and mutations. **r**, Violin plots of transition-frequency distributions for individual molecules, with dynamic modes and conditions indicated below; percentages above. **s-v**, Comparisons of the survival plots for each FRET state, fitted to tri- or bi-exponential functions across various Glu (**s**) or Asp (**t**) concentrations, CHS content (**u**), and mutations (**v**). **u-x**, Comparisons of the dwell time distribution for each FRET state across various Glu (**w**) or Asp (**x**) concentrations, CHS content (**y**), and mutations (**z**). Data are means of three independent repeats ± SEM.

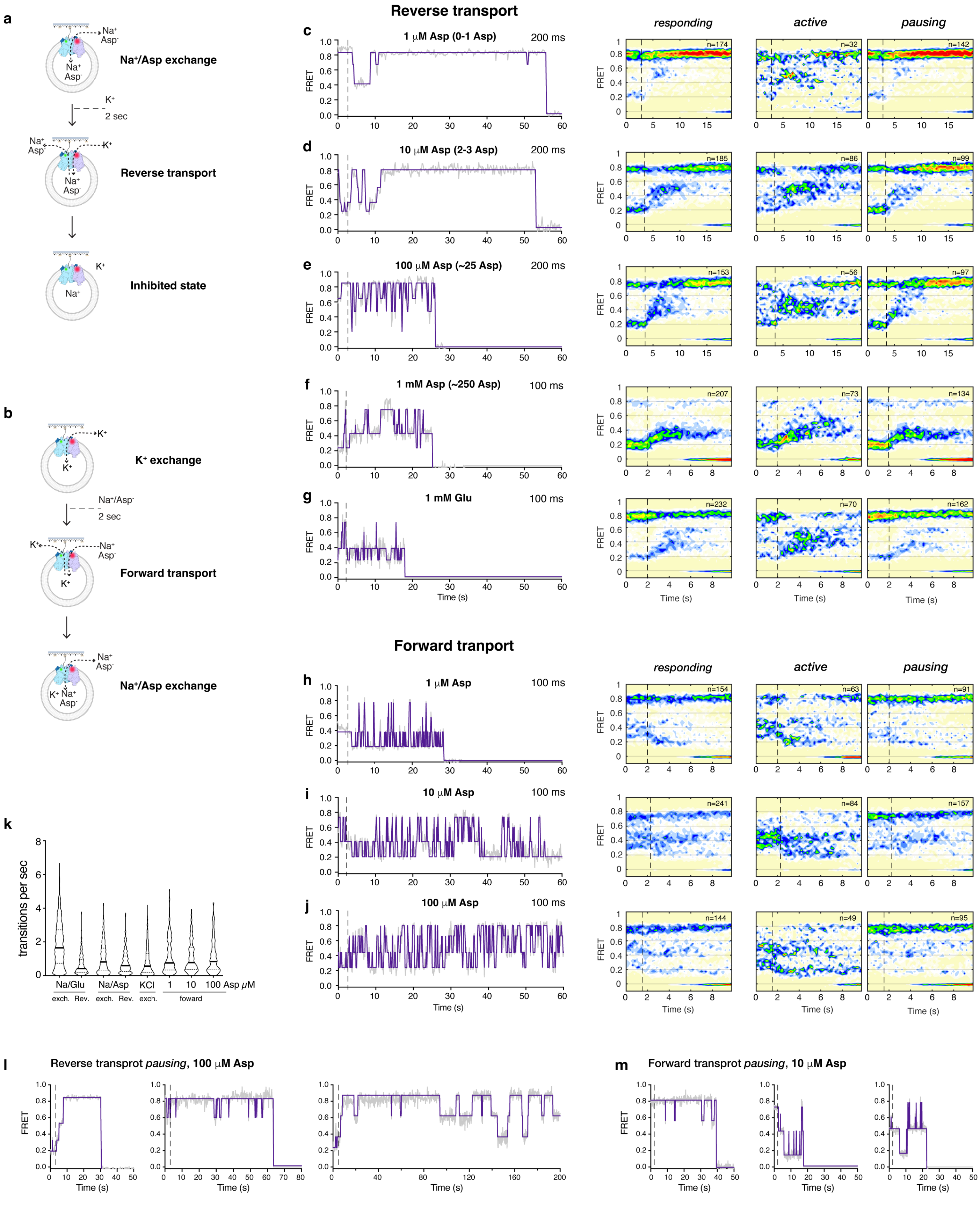


**Extended data figure 10. EAAT1$_{fret}$ dynamics under efflux and uptake transport conditions.**

**a–b**, Experimental scheme for efflux (reverse transport, **a**) and uptake (forward transport, **b**) activity measurements. WT EAAT1$_{fret}$ proteoliposomes were prepared in $K^+$ buffer or $Na^+$/Asp buffers with varying Asp concentrations. Recordings were initiated, and after ~2 seconds, proteoliposomes were perfused with $Na^+$/Asp (forward) or $K^+$ (reverse) buffer. **c-j**, From left to right: example smFRET traces and population plots for molecules that responded to perfusion ("*responded*"), including those that continued rounds of transport dynamics ("*active*") and those that exhibited pausing features or paused soon after reaching the ~0.8 FRET state ("*pausing*"), for reverse (**c-g**) and forward (**h–j**) transport. Raw data are in gray, and idealizations are in purple. Asp or Glu concentrations are indicated above the plots. Dotted lines mark perfusion; molecule numbers (*n*) are shown on the population plots. Transition frequency distributions for forward and reverse transport (**k**). Example smFRET traces exhibiting pausing features or paused soon after reaching the ~0.8 FRET state under reverse (**l**) and forward (**m**) transport conditions, respectively.

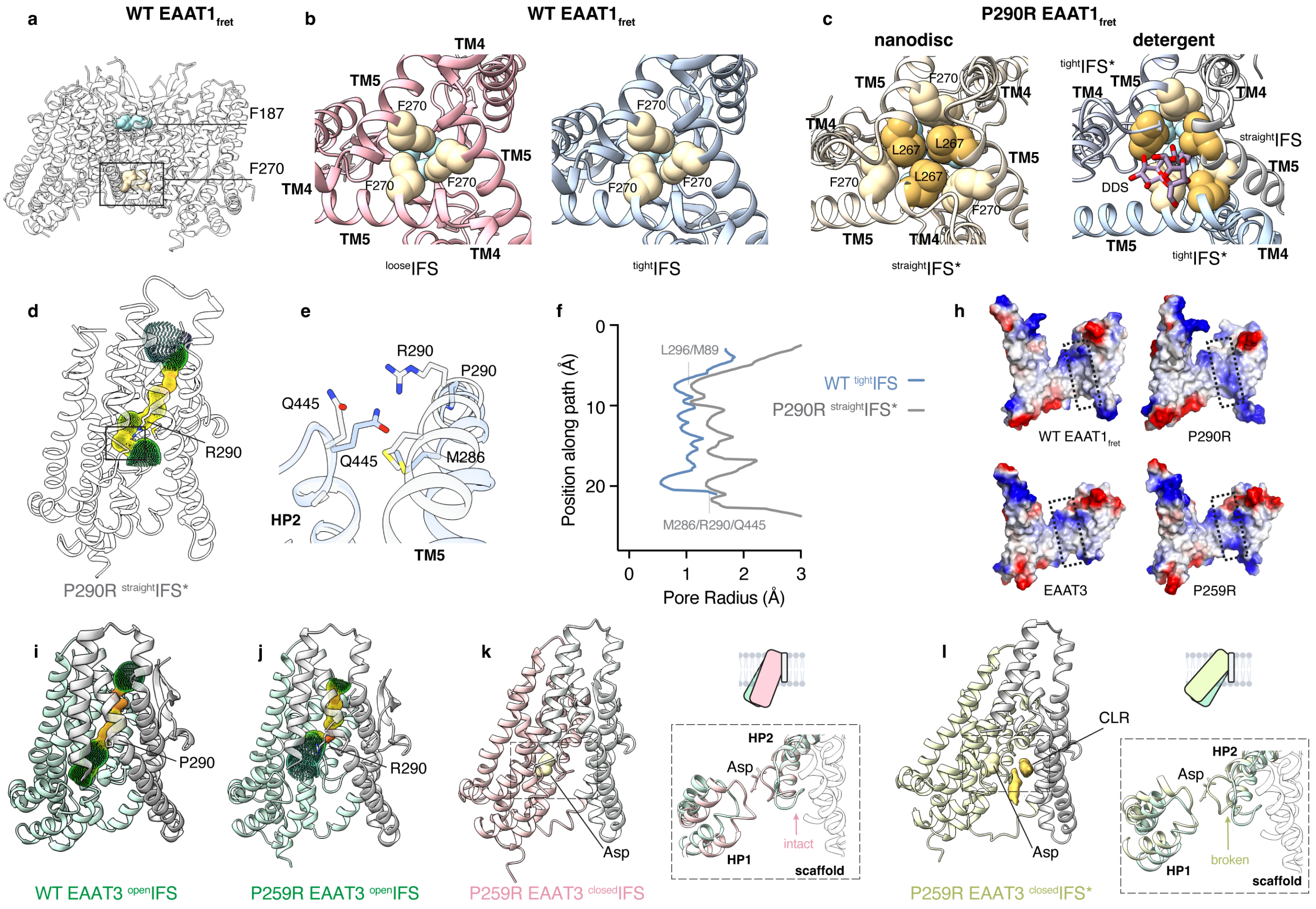


**Extended data figure 11. Structures of the P290R mutant of EAAT1$_{fret}$ and the P259R mutant of EAAT3.**
**a**, Cartoon of EAAT1$_{fret}$ in the IFS, viewed in the membrane plane. Phenylalanine clusters F187 (blue) and F270 (sand) at the extracellular and cytoplasmic sides of the trimer interface are shown as spheres. **b-c**, Cytoplasmic views of WT (**b**) and P290R (**c**). **b**, Trimers assembled from $^{loose}$IFS (pink) or $^{tight}$IFS (light blue) protomers. **c**, In nanodiscs, P290R forms a symmetric trimer of $^{straight}$IFS* protomers (left, light beige); in detergent, an asymmetric trimer (right) of $^{tight}$IFS (light blue), $^{tight}$IFS* (gray), and $^{straight}$IFS (white). In $^{tight}$IFS*, a DDS molecule (sticks, atom-colored) inserts into the trimer interface. **d**, $Cl^-$ conductance pore of a $^{straight}$IFS* protomer calculated with HOLE. Radius ranges from narrow (red) to wide (green); the black rectangle marks the intracellular constriction. 290R shown as sticks. **e**, Cytoplasmic view of the constriction. WT $^{tight}$IFS (blue) and P290R $^{straight}$IFS* (white) superimposed on the transport domain; constriction residues shown as sticks. **f**, Pore-radius profiles of WT $^{tight}$IFS (blue) and P290R $^{straight}$IFS* (gray). **h**, Scaffold domains shown as electrostatic surfaces for WT $^{tight}$IFS and P290R $^{straight}$IFS* (top) and WT EAAT3 $^{open}$IFS and P259R $^{open}$IFS (bottom). Dotted rectangles indicate pore-lining regions; colors range from blue (positive) to red (negative). **I-j**, $Cl^-$ pores (HOLE) in WT EAAT3 $^{open}$IFS (**i**) and P259R $^{open}$IFS (**j**). Scaffold domains, gray; transport domains, green; residue 259 shown as sticks. **k-l**, Two substrate-bound IFS classes from the P259R EAAT3 dataset: $^{closed}$IFS (**k**) and $^{CLR}$IFS (**l**), aligned to $^{open}$IFS on the scaffold domain. Transport domains colored green ($^{open}$IFS), pink ($^{closed}$IFS), and lime ($^{CLR}$IFS). Bound Asp shown as spheres; cholesterol density, dark yellow. Dotted lines indicate close-ups at right; schematics above show transport-domain movements. $^{closed}$IFS retains an intact HP2-scaffold interface; in $^{CLR}$IFS, the transport domain tilts away, with two cholesterol molecules occupying the interdomain space.

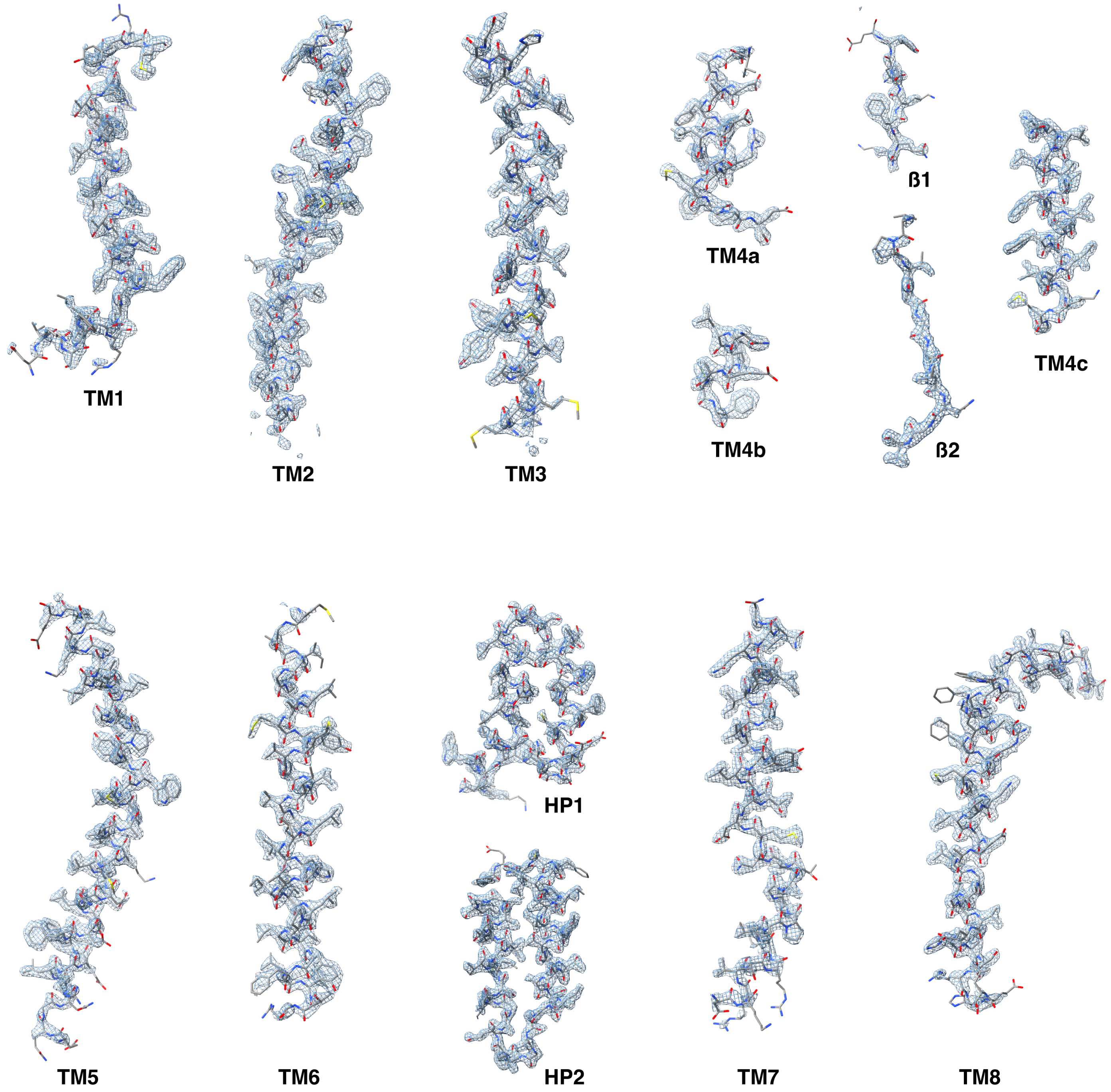


**Extended data figure 12. CryoEM density for TMs from $^{tight}$IFS.**
The cryoEM maps (shown as a mesh) for TMs 1-8, the β sheets 1-2, and the hairpins (HPs) 1-2, with the built model (shown in stick representation).

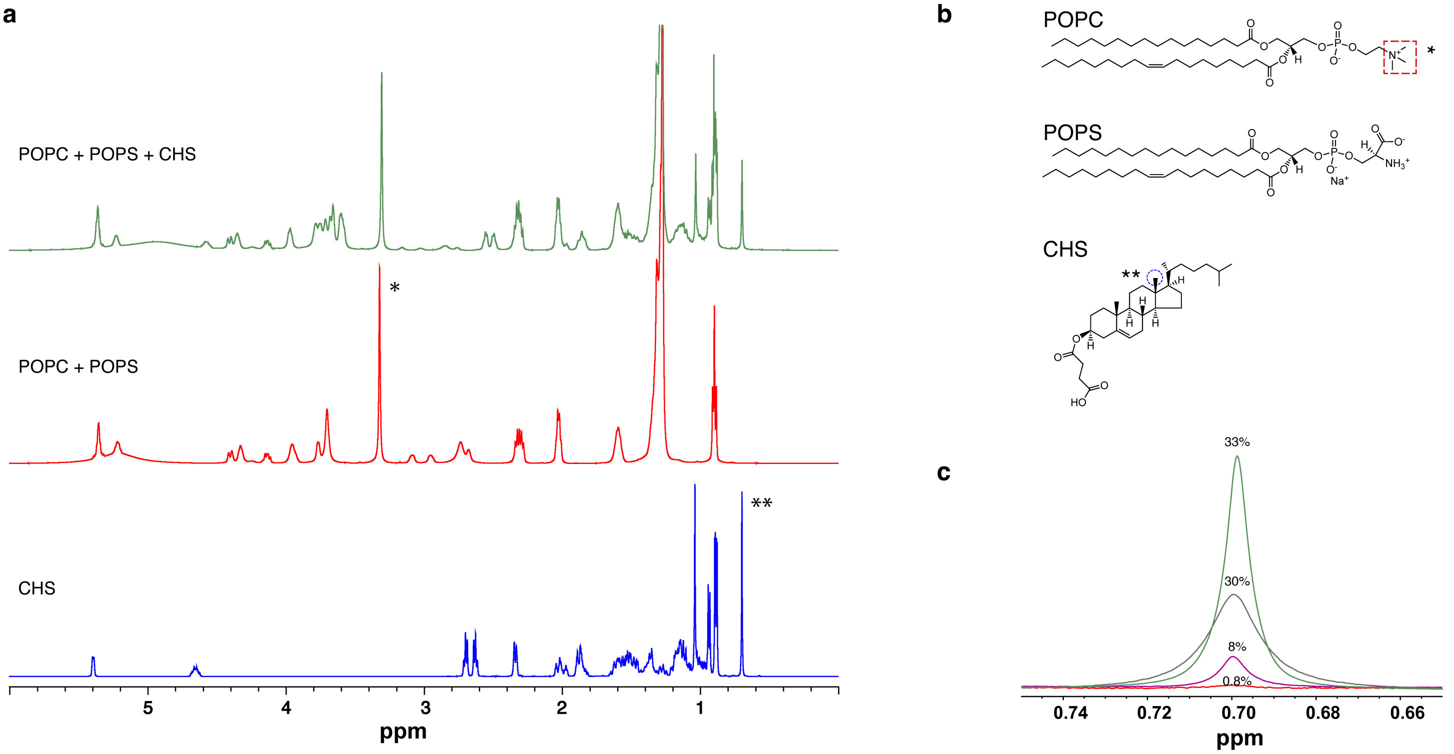


**Extended data figure 13. Cholesteryl hemisuccinate (CHS) content in liposomes measured by NMR**. **a**, Representative $^{1}$H-NMR spectra of an EAAT1$_{fret}$ proteoliposome preparation containing POPC and POPS, with (green) or without CHS (red), as well as pure CHS (blue). The -N($CH_3$)$_3$ peak in POPC (*) and the 18-$CH_3$ peak in CHS (**) are indicated. **b**, Chemical structures of POPC, POPS, and CHS. **c**, Zoomed-in view of the 18-$CH_3$ peaks of the stacked spectra of proteoliposome preparations, normalized to the intensity of the -N($CH_3$)$_3$ peaks. For each sample, the CHS fraction (percentages shown above the curves) is calculated as the ratio of the peak areas of 18-$CH_3$ and -N($CH_3$)$_3$.

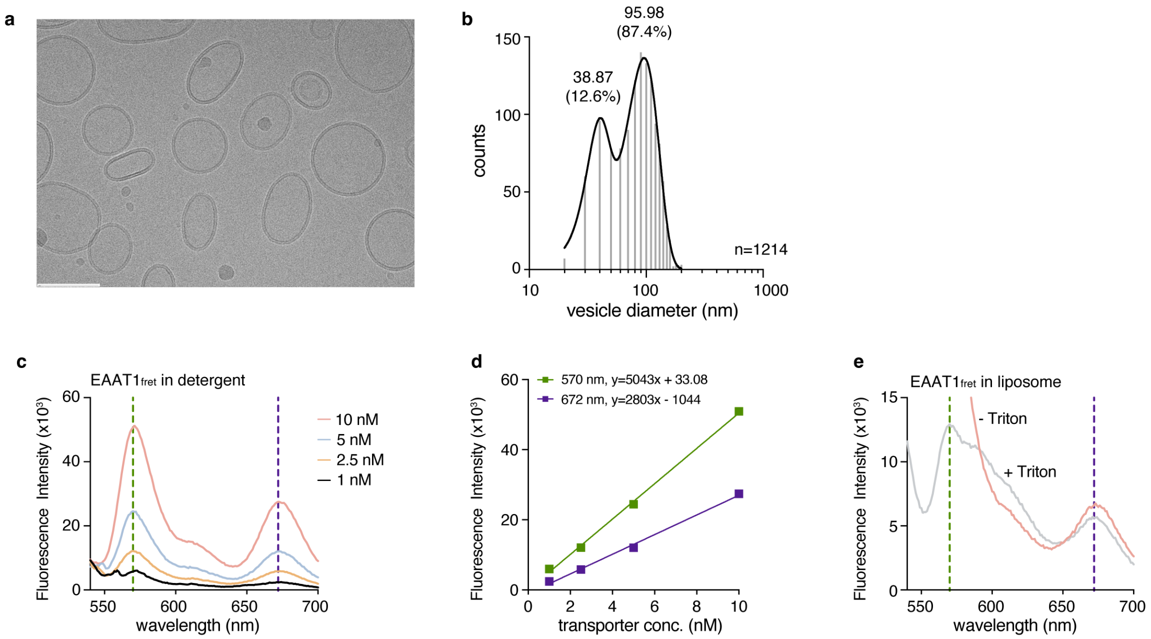


**Extended data figure 14. Proteoliposomes for the smFRET experiments. a,** A representative cryoEM micrograph of $EAAT1_{fret}$ proteoliposomes used in smFRET experiments. The scale bar (bottom left) indicates 100 nm. **b,** Histogram of vesicle diameters, calculated from ImageJ-measured circumferences and fitted with two Gaussians. The number of vesicles measured (n) is shown on the graph. The diameters in nm corresponding to the distribution peaks and the vesicle populations of each peak are indicated above the graph. **c-e**, Estimation of $EAAT1_{fret}$ reconstitution efficiency. Fluorescence emission spectra were recorded for detergent-solubilized labeled $EAAT1_{fret}$ at different concentrations; excitation wavelength was 532 nm (**c**). Emission intensities at 570 nm (LD555, green) and 672 nm (LD555-to-LD655-FRET, purple) were used to construct standard curves for estimating transporter concentration (**d**). Emission spectra of $EAAT1_{fret}$ proteoliposomes before (red) and after (gray) solubilization with Triton-X100 (**e**). Emission signals at 570 nm and 672 nm, and the standard curves from (**d**), were used to calculate reconstitution efficiencies of 32.75% and 32.87%, respectively. Representative data are shown.

Extended Data Table 1. Cryo-EM data collection, refinement, and validation statistics

| | $^{tight}$IFS$_{NaAsp}$ **9ZDT** **EMD-74073** | $^{loose}$IFS$_{NaAsp}$ **9ZDZ** **EMD-74079** | $^{loose}$IFS*$_{NaAsp}$ **9ZQK** **EMD-74568** | **V309G** $^{loose}$IFS$_{NaAsp}$ **9ZQL** **EMD-74569** | **V309G** $^{loose}$IFS*$_{NaAsp}$ **9ZQM** **EMD-74570** | **Y288L** $^{tight}$IFS$_{NaAsp}$ **9ZKB** **EMD-74360** |
|---|---|---|---|---|---|---|
| **Data collection and processing** | | | | | | |
| Magnification | 105,000 | | | 81,000 | | 105,000 |
| Voltage (kV) | 300 | | | 300 | | 300 |
| Electron exposure ($e^{-}$/Å$^2$) | 59.19 | | | 60 | | 58 |
| Defocus range (μm) | -0.9 to -1.9 | | | -1.0 to -2.3 | | -0.4 to -2.8 |
| Pixel size (Å) | 0.4125 | | | 0.426 | | 0.415 |
| Initial particle images (no.) | 4,663,691 | | | 7,405,793 | | 7,714,610 |
| Final particle images (no.) | 2,007,616 | | | 1,028,938 | | 681,355 |
| **Symmetry expansion (C3)** | | | | | | |
| Classified particle images (no.) | 1,158,774 | 1,120,942 | 794,852 | 528,980 | 516,598 | 502,063 |
| Map resolution (Å) FSC threshold **0.143** | 2.15 | 2.23 | 2.51 | 2.56 | 2.79 | 2.41 |
| Map resolution range (Å) | 1.8 – 3.6 | 1.8 – 4.6 | 1.8 – 5.6 | 1.8 – 4.3 | 1.8 – 4.7 | 1.8 – 4.7 |
| **Refinement** | | | | | | |
| Initial model used (PDB code) | 7NPW | 9ZDT | 9ZDZ | 9ZDZ | 9ZQK | 9ZDT |
| Model resolution (Å) FSC threshold **0.5** | 2.3 | 2.4 | 2.8 | 2.9 | 3.1 | 3.0 |
| Map sharpening *B* factor (Å$^2$) | -77.3 | -78.8 | -82.3 | -107.2 | -126.4 | -70.7 |
| Model composition | | | | | | |
| Non-hydrogen atoms | 3337 | 3342 | 3216 | 3399 | 3293 | 3334 |
| Protein residues | 430 | 422 | 420 | 429 | 425 | 430 |
| Ligands | NA: 3 / ASP:1 Y01: 1 | NA: 3 / ASP:1 A1C2I: 2 / Y01: 1 | NA: 3 / ASP:1 | NA: 3 / ASP:1 A1C2I: 2 / Y01: 1 | NA: 3 / ASP:1 Y01: 1 | NA: 3 / ASP:1 Y01: 1 |
| *B* factors (Å$^2$) | | | | | | |
| Protein | 72.55 | 83.89 | 86.68 | 109.37 | 108.79 | 78.53 |
| Ligand | 81.25 | 101.83 | 104.02 | 133.33 | 120.59 | 94.01 |
| R.m.s. deviations | | | | | | |
| Bond lengths (Å) | 0.004 | 0.004 | 0.004 | 0.004 | 0.004 | 0.004 |
| Bond angles (°) | 0.944 | 0.921 | 0.962 | 0.955 | 0.910 | 0.966 |
| Validation | | | | | | |
| MolProbity score | 1.05 | 1.02 | 1.12 | 1.06 | 1.09 | 1.07 |
| Clashscore | 2.62 | 2.35 | 2.88 | 2.75 | 2.95 | 2.77 |
| Poor rotamers (%) | 0.00 | 0.00 | 0.00 | 0.28 | 0.00 | 0.28 |
| Ramachandran plot | | | | | | |
| Favored (%) | 98.11 | 99.04 | 97.83 | 99.05 | 99.52 | 99.06 |
| Allowed (%) | 1.89 | 0.96 | 2.17 | 0.95 | 0.48 | 0.94 |
| Disallowed (%) | 0.00 | 0.00 | 0.00 | 0.00 | 0.00 | 0.00 |

#### Extended Data Table 1. Cryo-EM data collection, refinement, and validation statistics (continued)

| | OFS$_{Apo}$<br>9ZQH<br>EMD-74565 | IFS$_{Apo}$<br>9ZQJ<br>EMD-74567 | $^{free}$OFS$_{NaAsp}$<br>9ZQW<br>EMD-74583 | $^{wrapped}$OFS$_{NaAsp}$<br>9ZR8<br>EMD-74596 | intS$_{NaAsp}$<br>9ZTT<br>EMD-74772 | $^{shift}$IFS$_{NaAsp}$<br>9ZU2<br>EMD-74788 | TFB-TBOA<br>$^{loose}$IFS$_{NaTFB}$<br>9ZU9<br>EMD-74799 | TFB-TBOA<br>$^{loose}$IFS*$_{NaTFB}$<br>9ZUE<br>EMD-74808 |
|---|---|---|---|---|---|---|---|---|
| **Data collection and processing** | | | | | | | | |
| Magnification | 105,000 | | 105,000 | | | 105,000 | 105,000 | |
| Voltage (kV) | 300 | | 300 | | | 300 | 300 | |
| Electron exposure (e$^{-}$/Å$^{2}$) | 49.97 | | 51.88 | | | 47.77 | 57.98 | |
| Defocus range (μm) | -0.6 to -2.5 | | -0.4 to -2.6 | | | -0.6 to -2.5 | -0.7 to -2.2 | |
| Pixel size (Å) | 0.4125 | | 0.4125 | | | 0.4125 | 0.4125 | |
| Initial particle images (no.) | 1,851,475 | | 1,774,159 | | | 2,487,687 | 1,769,001 | |
| Final particle images (no.) | 326,891 | | 661,346 | | | 490,349 | 920,551 | |
| **Symmetry expansion (C3)** | | | | | | | | |
| Classified particle images (no.) | 224,439 | 195,005 | 130,827 | 121,145 | 110,183 | 170,449 | 585,400 | 424,063 |
| Map resolution (Å)<br>FSC threshold **0.143** | 2.61 | 3.01 | 2.94 | 2.93 | 3.18 | 3.05 | 2.60 | 2.77 |
| Map resolution range (Å) | 1.8 – 4.3 | 2.0 – 5.8 | 1.8 – 6.7 | 1.9 – 8.3 | 2.1 – 9.4 | 1.9 – 7.0 | 1.8 – 5.5 | 1.8 – 5.6 |
| **Refinement** | | | | | | | | |
| Initial model used (PDB code) | 9ZQW | 7NPW | 5LLU | 9ZQW | 9ZQW | 9ZDZ | 9ZQJ | 9ZQJ |
| Model resolution (Å)<br>FSC threshold **0.5** | 2.8 | 3.3 | 3.2 | 3.1 | 3.6 | 3.3 | 2.9 | 3.1 |
| Map sharpening *B* factor (Å$^{2}$) | -135.0 | -98.6 | -98.8 | -99.5 | -107.7 | -91.7 | -93.2 | -96.6 |
| Model composition | | | | | | | | |
| Non-hydrogen atoms | 3230 | 3192 | 3200 | 3306 | 3223 | 3205 | 3211 | 3180 |
| Protein residues | 424 | 415 | 419 | 424 | 422 | 415 | 410 | 410 |
| Ligands | 0 | Y01: 1 | NA: 3 / ASP: 1 | NA: 3 / ASP: 1<br>Y01: 2 | NA: 3 / ASP: 1 | NA: 2 / ASP: 1<br>Y01: 1 | NA: 2 / 7O9:1<br>A1C2I: 1 /<br>Y01: 1 | NA: 2 / 7O9:1<br>Y01: 1 |
| *B* factors (Å$^{2}$) | | | | | | | | |
| Protein | 91.82 | 119.09 | 84.98 | 97.04 | 31.29 | 96.06 | 79.45 | 81.33 |
| Ligand | - | 149.19 | 102.89 | 110.62 | 39.29 | 113.89 | 101.88 | 105.73 |
| R.m.s. deviations | | | | | | | | |
| Bond lengths (Å) | 0.004 | 0.004 | 0.004 | 0.004 | 0.004 | 0.004 | 0.004 | 0.004 |
| Bond angles (°) | 0.922 | 0.960 | 0.936 | 0.919 | 0.990 | 0.927 | 0.994 | 0.911 |
| Validation | | | | | | | | |
| MolProbity score | 0.98 | 1.30 | 1.03 | 1.11 | 1.35 | 1.09 | 1.04 | 1.06 |
| Clashscore | 2.10 | 5.47 | 2.43 | 3.22 | 6.33 | 3.03 | 2.59 | 2.75 |
| Poor rotamers (%) | 0.00 | 0.00 | 0.00 | 0.00 | 0.00 | 0.00 | 0.00 | 0.00 |
| Ramachandran plot | | | | | | | | |
| Favored (%) | 99.04 | 98.29 | 99.52 | 99.29 | 98.33 | 98.78 | 99.01 | 98.76 |
| Allowed (%) | 0.96 | 1.71 | 0.48 | 0.71 | 1.67 | 1.22 | 0.99 | 1.24 |
| Disallowed (%) | 0.00 | 0.00 | 0.00 | 0.00 | 0.00 | 0.00 | 0.00 | 0.00 |

**Extended Data Table 1. Cryo-EM data collection, refinement, and validation statistics** (continued)

| | P290R $^{straight}IFS_{NaAsp}$ 10BN EMD-75046 | P290R $^{tight}IFS_{NaAsp}$ 10BQ EMD-75051 | P290R $^{tight}IFS*_{NaAsp}$ 10BS EMD-75052 | P290R $^{straight}IFS*_{NaAsp}$ 10BP EMD-75050 | P259R E3 $^{open}IFS_{Na}$ 10CI EMD-75063 | P259R E3 $^{closed}IFS_{NaAsp}$ 10CJ EMD-75064 | P259R E3 $^{closed}IFS*_{NaAsp}$ 10CO EMD-75069 |
|---|---|---|---|---|---|---|---|
| **Data collection and processing** | | | | | | | |
| Magnification | | 105,000 | | 105,000 | | 105,000 | |
| Voltage (kV) | | 300 | | 300 | | 300 | |
| Electron exposure (e⁻/Å²) | | 52.58 | | 48.6 | | 51.1 | |
| Defocus range (μm) | | -0.8 to -2.3 | | -0.7 to -3.2 | | -1.1 to -2.0 | |
| Pixel size (Å) | | 0.4125 | | 0.4125 | | 0.426 | |
| Initial particle images (no.) | | 4,000,305 | | 1,382,819 | | 3,694,569 | |
| Final particle images (no.) | | 1,559,306 | | 108,251 | | 154,187 | |
| **Symmetry expansion (C3)** | | | | | | | |
| Classified particle images (no.) | 1,006,502 | 685,402 | 747,478 | 83,474 | 125,506 | 118,207 | 117,205 |
| Map resolution (Å) FSC threshold **0.143** | 2.77 | 2.81 | 2.77 | 3.65 | 2.92 | 3.10 | 3.04 |
| Map resolution range (Å) | 1.8 – 6.2 | 1.8 – 5.2 | 1.8 – 6.2 | 2.4 – 7.6 | 2.0 – 4.7 | 1.9 – 5.9 | 2.0 – 26.3 |
| **Refinement** | | | | | | | |
| Initial model used (PDB code) | 9ZDT | 9ZDT | 10BQ | 10BN | 9P4Z | 10CI | 10CJ |
| Model resolution (Å) FSC threshold **0.5** | 3.0 | 3.0 | 3.0 | 4.0 | 3.3 | 3.4 | 3.5 |
| Map sharpening *B* factor (Å²) | -125.8 | -127.9 | -129.6 | -107.7 | -82.1 | -91.1 | -85.2 |
| Model composition | | | | | | | |
| Non-hydrogen atoms | 3210 | 3188 | 3345 | 3144 | 3120 | 3086 | 3035 |
| Protein residues | 415 | 417 | 423 | 412 | 409 | 404 | 392 |
| Ligands | NA: 3 / ASP:1 / Y01: 1 | NA: 3 / ASP: 1 | NA: 3 / ASP:1 A1C2I: 1 / Y01: 2 | ASP:1 | NA: 1 | ASP: 1 / NA:3 | ASP: 1 / CLR: 2 |
| *B* factors (Å²) | | | | | | | |
| Protein | 105.03 | 106.09 | 100.26 | 26.96 | 88.99 | 102.10 | 64.07 |
| Ligand | 119.41 | 60.83 | 114.10 | 22.95 | 97.62 | 102.91 | 19.53 |
| R.m.s. deviations | | | | | | | |
| Bond lengths (Å) | 0.004 | 0.004 | 0.004 | 0.004 | 0.004 | 0.004 | 0.004 |
| Bond angles (°) | 0.945 | 0.936 | 0.907 | 0.923 | 0.934 | 0.940 | 0.931 |
| Validation | | | | | | | |
| MolProbity score | 1.15 | 1.25 | 1.18 | 1.28 | 1.10 | 1.21 | 1.23 |
| Clashscore | 3.63 | 4.88 | 3.94 | 5.25 | 3.11 | 4.25 | 4.29 |
| Poor rotamers (%) | 0.00 | 0.00 | 0.00 | 0.00 | 0.00 | 0.00 | 0.00 |
| Ramachandran plot | | | | | | | |
| Favored (%) | 99.51 | 99.03 | 98.32 | 98.51 | 98.26 | 98.74 | 97.92 |
| Allowed (%) | 0.49 | 0.97 | 1.68 | 1.49 | 1.74 | 1.26 | 2.08 |
| Disallowed (%) | 0.00 | 0.00 | 0.00 | 0.00 | 0.00 | 0.00 | 0.00 |

**Extended Data Table 2.** Distances, expected FRET states, and schematic representation of states.

| protomer 1 | protomer 2 | distance | estimated FRET | measured FRET | |
|---|---|---|---|---|---|
| $^{wrapped}$OFS$_{NaAsp}$ | $^{wrapped}$OFS$_{NaAsp}$ | 81.6 | 0.16 | | |
| $^{free}$OFS$_{NaAsp}$ | $^{free}$OFS$_{NaAsp}$ | 78.7 | 0.19 | | 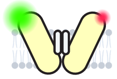 |
| $^{wrapped}$OFS$_{NaAsp}$ | $^{free}$OFS$_{NaAsp}$ | 80 | 0.17 | NaAsp, 0.21 | |
| $^{free}$OFS$_{Apo}$ | $^{free}$OFS$_{Apo}$ | 76.5 | 0.22 | NaCl, 0.23 | |
| $^{wrapped}$OFS$_{NaAsp}$ | intS | 73.5 | 0.27 | KCl, 0.26 | |
| $^{free}$OFS$_{NaAsp}$ | intS | 72.2 | 0.29 | | 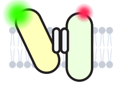 |
| $^{free}$OFS$_{Apo}$ | intS | 71.5 | 0.30 | | |
| intS | intS | 65.4 | 0.42 | | |
| $^{loose}$IFS$_{NaAsp}$ | $^{wrapped}$OFS$_{NaAsp}$ | 64.9 | 0.43 | | |
| $^{tight}$IFS$_{NaAsp}$ | $^{wrapped}$OFS$_{NaAsp}$ | 64.5 | 0.44 | | |
| $^{loose}$IFS$_{NaAsp}$ | $^{free}$OFS$_{NaAsp}$ | 62.9 | 0.48 | | 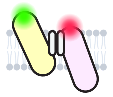 |
| $^{tight}$IFS$_{NaAsp}$ | $^{free}$OFS$_{NaAsp}$ | 62.6 | 0.49 | | |
| $^{wrapped}$OFS$_{NaAsp}$ | $^{int}$IFS$_{NaAsp}$ | 66.4 | 0.40 | NaAsp, 0.42 | |
| $^{free}$OFS$_{NaAsp}$ | $^{int}$IFS-NaAsp | 65.0 | 0.43 | NaCl, 0.44 | |
| $^{loose}$IFS$_{Na}$ | $^{wrapped}$OFS$_{NaAsp}$ | 64.3 | 0.45 | KCl, 0.47 | |
| $^{preApo}$IFS$_{Na}$ | $^{wrapped}$OFS$_{NaAsp}$ | 64.5 | 0.44 | | 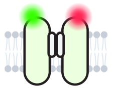 |
| IFS$_{Apo}$ | $^{wrapped}$OFS$_{NaAsp}$ | 64.4 | 0.44 | | |
| IFS$_{Apo}$ | $^{free}$OFS$_{NaAsp}$ | 62.6 | 0.49 | | |
| $^{loose}$IFS$_{Na}$ | $^{free}$OFS$_{NaAsp}$ | 62.4 | 0.49 | | |
| $^{preApo}$IFS$_{Na}$ | $^{free}$OFS$_{NaAsp}$ | 62.6 | 0.48 | | |
| $^{loose}$IFS$_{NaAsp}$ | $^{free}$OFS$_{Apo}$ | 62.2 | 0.50 | | |
| $^{tight}$IFS$_{NaAsp}$ | $^{free}$OFS$_{Apo}$ | 61.9 | 0.50 | | |
| $^{loose}$IFS$_{Na}$ | $^{free}$OFS$_{Apo}$ | 61.7 | 0.51 | | |
| $^{preApo}$IFS$_{Na}$ | $^{free}$OFS$_{Apo}$ | 61.9 | 0.50 | | |
| IFS$_{Apo}$ | $^{free}$OFS$_{Apo}$ | 61.8 | 0.50 | | |
| $^{loose}$IFS$_{NaAsp}$ | intS | 56.0 | 0.65 | | |
| $^{tight}$IFS$_{NaAsp}$ | intS | 55.6 | 0.66 | NaAsp, 0.59 | 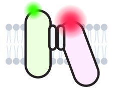 |
| $^{loose}$IFS$_{Na}$ | intS | 55.4 | 0.66 | NaCl, 0.60 | |
| $^{preApo}$IFS$_{Na}$ | intS | 55.7 | 0.66 | KCl, 0.61 | |
| IFS$_{Apo}$ | intS | 55.6 | 0.66 | | |
| $^{int}$IFS$_{NaAsp}$ | intS | 57.8 | 0.60 | | |
| $^{tight}$IFS$_{NaAsp}$ | $^{tight}$IFS$_{NaAsp}$ | 45.2 | 0.87 | | |
| $^{loose}$IFS$_{NaAsp}$ | $^{loose}$IFS$_{NaAsp}$ | 45.6 | 0.86 | | |
| $^{tight}$IFS$_{NaAsp}$ | $^{loose}$IFS$_{NaAsp}$ | 45.4 | 0.87 | | |
| $^{loose}$IFS$_{Na}$ | $^{loose}$IFS$_{Na}$ | 44.3 | 0.88 | | |
| $^{loose}$IFS$_{Na}$ | $^{preApo}$IFS$_{Na}$ | 44.7 | 0.88 | NaAsp, 0.78 | 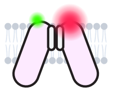 |
| $^{preApo}$IFS$_{Na}$ | $^{preApo}$IFS$_{Na}$ | 44.8 | 0.87 | NaCl, 0.78 | |
| IFS$_{Apo}$ | IFS$_{Apo}$ | 45.2 | 0.87 | KCl, 0.79 | |
| $^{int}$IFS$_{NaAsp}$ | $^{int}$IFS$_{NaAsp}$ | 49.6 | 0.79 | | |
| $^{loose}$IFS$_{NaAsp}$ | $^{int}$IFS$_{NaAsp}$ | 48.0 | 0.82 | | |
| $^{tight}$IFS$_{NaAsp}$ | $^{int}$IFS$_{NaAsp}$ | 47.7 | 0.83 | | |
| $^{loose}$IFS$_{Na}$ | $^{int}$IFS$_{NaAsp}$ | 47.4 | 0.83 | | |
| $^{preApo}$IFS$_{Na}$ | $^{int}$IFS$_{NaAsp}$ | 47.7 | 0.83 | | |
| IFS$_{Apo}$ | $^{int}$IFS$_{NaAsp}$ | 47.6 | 0.83 | | |
